# Distribution of big tau isoforms in the human central and peripheral nervous system

**DOI:** 10.1101/2025.09.15.676127

**Authors:** Rama Krishna Koppisetti, Nicolas R. Barthélemy, Kanta Horie, Cindy V. Ly, Kaleigh F. Roberts, Srinivas Koutarapu, Richard J. Perrin, Erin E. Franklin, Chiara Pedicone, Joshua Orrick, Justin Melendez, Timothy M. Miller, Chihiro Sato, Nupur Ghoshal, Alison M. Goate, Celeste M. Karch, Randall J. Bateman, Soumya Mukherjee

## Abstract

**Objective:** Tau is widely studied in the field of neurodegenerative disease research, yet most work has focused on canonical brain tau isoforms. A longer isoform, “big tau,” produced by inclusion of exon 4a, is expressed in the peripheral nervous system (PNS) and select central nervous system (CNS) regions. We sought to characterize big tau molecular composition, anatomical distribution, and relevance to neurodegenerative disease.

**Methods:** Mass spectrometry was used to sequence big tau and map its distribution across the human nervous system. Postmortem samples included brain tissue from Alzheimer’s disease (AD), amyotrophic lateral sclerosis (ALS), and controls; spinal cord from ALS and controls; and peripheral nerves. Big and canonical (“small”) tau isoforms were also quantified in cerebrospinal fluid (CSF) from young controls and participants stratified by amyloid status and cognitive impairment.

**Results:** Human big tau results from insertion of either 355 or 251 amino acids encoded by exon 4a-long and exon 4a-short, respectively. Alternative splicing of exons 2, 3, and 10 generates multiple big tau isoforms. Total tau levels were ∼1000-fold higher in brain than in the PNS; however, the relative abundance of big tau increased from the CNS to the PNS, comprising 50 % of the total tau in the periphery and exhibiting considerable regional heterogeneity in the brain (∼ 1 % of total tau). In CSF, big tau levels were unchanged by amyloid abnormalities or cognitive impairment, whereas canonical tau increased with AD-related pathology.

**Interpretation:** Big tau represents a distinct tau population enriched in the PNS and largely uncoupled from disease-associated changes in brain-derived tau, suggesting that distinguishing big tau from canonical tau may improve interpretation of tau biomarkers and help differentiate CNS neurodegeneration from peripheral nerve pathology.

## Introduction

Neurofibrillary tangles (NFTs) composed of microtubule associated protein tau (MAPT) are one of the key pathological hallmarks of Alzheimer’s disease (AD) along with extracellular amyloid-β plaques.^1–8^ Tau pathology within neocortical brain regions is closely linked to cognitive impairment in AD.^9–12^ Central nervous system (CNS)-specific tau phosphorylation (p-tau) have been validated as reliable indicators of CNS tau changes in response to AD pathology.^1^ The presence of AD neuropathological changes in the brain often coincides with an increase in tau and p-tau species in the cerebrospinal fluid (CSF), making CSF tau biomarkers good candidates for monitoring amyloidosis, NFT formation and cognitive decline in AD.^13–15^ Recent development of blood-based tau biomarkers has highlighted that tau measured in plasma reflects at least two sources: brain-derived tau released from the CNS and tau originating from peripheral tissues. ^16–18^ Peripheral tau species therefore comprise a substantial fraction of the total plasma tau pool, complicating the direct interpretation of blood tau measurements as Alzheimer’s disease–specific biomarkers. Discriminating CNS-derived tau isoforms from those originating in the peripheral nervous system (PNS) may improve the specificity of tau-based biomarkers for AD and other CNS neurodegenerative disorders. ^19,20^ Conversely, PNS-enriched tau isoforms could provide biomarkers for neurological diseases affecting the peripheral nervous system.^21–23^ However, our understanding of peripheral nervous system (PNS)-derived tau in humans remains limited. Characterizing the distribution of PNS-specific tau isoforms alongside the canonical CNS tau isoforms (referred to as “small tau” henceforth) will be critical for understanding tau biology and its contribution to the pathophysiology of neurodegenerative disorders in which tau is implicated.

Tau is encoded by the *MAPT* gene located on chromosome 17 in humans^24^, contains 16 exons^25^ and displays remarkable length heterogeneity in its isoforms due to alternative splicing of exons 2 and 3 at the N terminus and exon 10 in the microtubule binding region (MTBR)^26^. This leads to the generation of six major isoforms in the human CNS; 0N3R, 1N3R, 2N3R, 0N4R, 1N4R and 2N4R. Expression of these isoforms is developmentally regulated,^27,28^ with 0N3R being mainly expressed at the fetal stage; then, approximately equivalent expression of 0N3R, 1N3R, 0N4R and 1N4R is maintained in the adult brain, where 2N is less abundant.^29^ The functional roles of individual tau isoforms remain an active area of investigation, in part because disruption of the normal 3R:4R tau isoform ratio in the brain is a hallmark of several tauopathies.^30^ Additionally, multiple post-translational modifications (PTMs) and truncations lead to a multitude of tau proteoforms known to regulate tau physiological functions and contribute to pathological conformations.^31–37^ Recent advances in Cryo-EM, mass spectrometry and genome sequencing techniques have demonstrated the heterogeneity of tau isoforms that are involved in various tauopathies, suggesting that splicing is of key importance in the neuropathological process.^32–34,38^ Alternative splicing of the *MAPT* gene produces three transcripts of 2, 6 and 9 kb that are differentially expressed in the nervous system^39,40^. Tau is also expressed in peripheral tissue (e.g. peripheral nerves, heart, skeletal muscle, kidney) outside the CNS, where it plays a vital role in metabolism, microtubule formation, and microtubule stabilization.^41–46^ While the 2 kb transcript produces tau for the cell nucleus, 6 kb transcript expression leads to the common six tau isoforms of 45-60 kDa,^2^ whereas the expression of the 9 kb mRNA results in a higher molecular weight isoform (110 kDa) that was originally cloned and sequenced from rat complementary DNA (cDNA) and named big tau.^39,47,48^

Big tau results from an exon insertion (exon 4a) between exons 4 and 5 in tau transcripts. Translation of this exon 4a dramatically increases the length of the projection domain relative to the canonical small tau isoforms.^39,45,46,49–51^ Early work in rodents demonstrated that big tau expression is not limited to the PNS; it also occurs in selective regions in spinal cord and brain-stem.^47,52^ Big tau expression in the superior cervical ganglion (SCG) of the autonomic nervous system, and in the trigeminal ganglion and dorsal root ganglia (DRG) of the PNS was found to be developmentally regulated, beginning late in the embryonic stage, and increasing postnatally.^40,53^ Investigations using big tau specific antibodies in rodents have documented the expression of exon 4a in the spinal motor neurons, retinal ganglion cells (RGC), optic nerves, the cerebellum and CNS neurons that extend processes into the periphery, including cranial nerve motor nuclei.^40,54^ The functional rationale for the transition to big tau from the CNS isoform of tau in specific neuronal populations remains undefined, as well as the pathological significance of the additional exon 4a in big tau.^55^

The big tau sequence in humans could result from the insertion of exon 4a in the tau transcript,^45,46,50^ however proteomic evidence has not been established. In this study, we investigated the molecular compositions and anatomical distributions of human big tau isoforms across the nervous system. Here, we provide the first mass spectrometry–based evidence for the translation and expression of the unique tau insertion domain encoded by exon 4a within the N-terminal projection region between exons 4 and 5, expanding the known repertoire of tau proteins expressed in the human nervous system. To define the distribution of big tau, we developed an immunoprecipitation–mass spectrometry (IP-MS) strategy to sequence and quantify big tau–specific peptides alongside canonical tau peptides across human nervous system tissues. Finally, to assess the relevance of big tau with aging and neurodegeneration, we quantified extracellular big tau peptides in CSF from individuals with and without AD. Together, these findings define the molecular diversity and anatomical distribution of big tau in humans and suggest that distinguishing big tau from canonical tau may enable improved biomarker strategies to differentiate CNS and PNS neurological diseases.

## Materials and Methods

### Human tissue samples and CSF

Frozen postmortem brain tissue samples from cases representing different stages of Alzheimer’s disease neuropathologic change (ADNC) were obtained from the Charles F. and Joanne Knight ADRC Neuropathology Core at Washington University School of Medicine.^56^ The studies involving postmortem human brain samples were approved by the Washington University Human Research Protection Office (HASD 201105103, ADRC – 201105102, ACS – 201105305). The brain regions studied included cerebellum (CB), superior frontal gyrus (SFG), superior temporal gyrus (STG), occipital pole (Occ) and inferior parietal lobule (P). This study was approved by the Washington University in St. Louis Institutional Review Board. Brain, spinal cord, and sciatic nerve samples from neuropathologically confirmed sporadic amyotrophic lateral sclerosis (ALS) and disease controls were obtained from the Washington University ALS Postmortem Core. Postmortem nervous tissues across the spinal cord regions and PNS (cauda equina, dorsal root ganglion, sciatic nerve and brachial plexus) were obtained from two healthy young control donors and stored at -80 . All participants provided consent for autopsy and research participation. The demographics of the frozen brain (Charles F. and Joanne Knight ADRC) cohort were previously described and shown in Table 1a.^56^ The brain, spinal cord and PNS sample demographics are also included in Table 1b.

**Table 1a:**
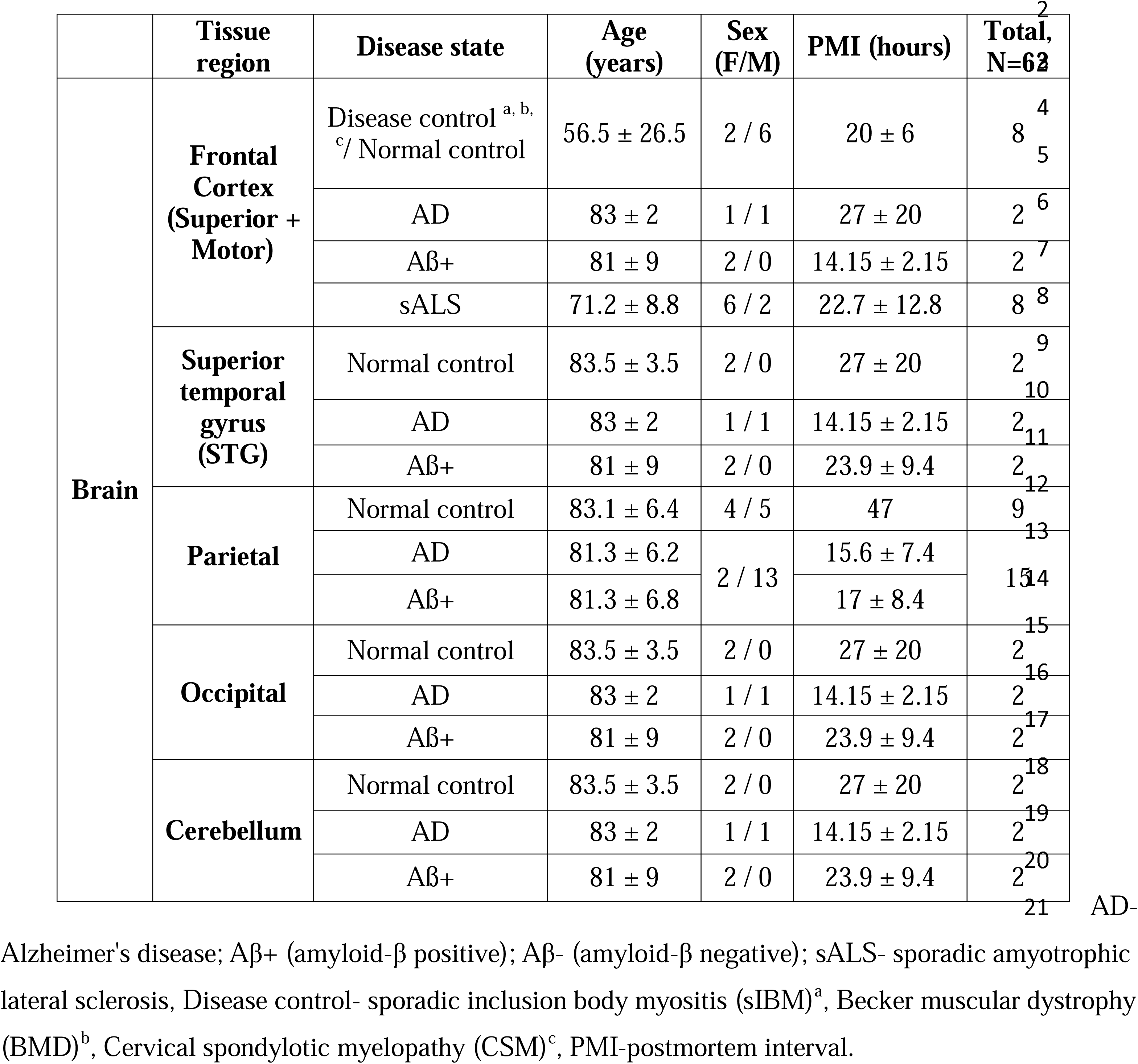
Demographics of the human brain samples.

**Table 1b.** Demographics of the human spinal cord, sciatic nerve, cauda equina, brachial plexus and dorsal root ganglion samples.

|  | Tissue region | Disease state | Age (years) | Sex (F/M) | PMI (hours) | Total, N=35 |
| --- | --- | --- | --- | --- | --- | --- |
| Spinal cord | Lumbar | Disease control <sup>a, b</sup> / Normal control | 48.1 ± 26.9 | 1 / 2 | 26 | 3 |
|  |  | sALS | 71.2 ± 8.8 | 3 / 1 | 22.7 ± 12.8 | 4 |
|  |  | Normal control | 40 ± 2 | 2 / 0 | 26 ± 3 | 2 |
|  | Cervical | Disease control <sup>a, b, c</sup> / Normal control | 56.5 ± 26.5 | 1 / 3 | 20 ± 6 | 4 |
|  |  | sALS | 71.2 ± 8.8 | 3 / 1 | 22.7 ± 12.8 | 4 |
|  |  | Normal control | 40 ± 2 | 2 / 0 | 26 ± 3 | 2 |
|  | Thoracic | Normal control | 38 | 1 / 0 | 29 | 1 |
|  | Sacral | Normal control | 38 | 1 / 0 | 29 | 1 |
| Sciatic nerve | Nerve | Disease control <sup>c</sup> | 81 | 0 / 1 | 14 | 1 |
|  |  | sALS | 63.6 ± 3.3 | 1 / 4 | 40.1 ± 8.3 | 5 |
|  |  | Normal control | 40 ± 2 | 2 / 0 | 26 ± 3 | 2 |
| Cauda equina | Nerve | Normal control |  | 2 / 0 |  | 2 |
| Brachial plexus | Nerve | Normal control |  | 2 / 0 |  | 2 |
| Dorsal root ganglion | Nerve | Normal control |  | 2 / 0 |  | 2 |

AD- Alzheimer's disease; Aβ <sup>+</sup> (amyloid-β positive); Aβ <sup>-</sup> (amyloid-β negative); sALS- sporadic amyotrophic lateral sclerosis, Disease control- sporadic inclusion body myositis (sIBM)<sup>a</sup>, Becker muscular dystrophy (BMD)<sup>b</sup>, Cervical spondylotic myelopathy (CSM)<sup>c</sup>, PMI-postmortem interval.

CSF samples were collected from 90 participants who underwent human tau Stable Isotope Labeling Kinetics (SILK) protocol as described previously.^57^ The research with CSF samples was approved by the Human Studies Committee and the General Clinical Research Center (GCRC Tau SILK: 201502091). Clinical and cognitive assessments, including the Clinical Dementia Rating (CDR^®^) and Mini-Mental State Examination (MMSE) scores at baseline CSF collection for the participants, were used in this study. CSF amyloid-β 42/40 ratio along with CDR scores were used for clinical group classification as follows: amyloid negative, cognitively unimpaired (CDR = 0); amyloid positive, cognitively unimpaired (CDR = 0, preclinical AD); and mild to moderate AD dementia (CDR = 0.5-2).

Amyloid-β 42/40 ratio was estimated using the IP-MS assay and amyloid status was defined using amyloid-β 42/40 ratio cutoff scores as previously reported.^58^ The corresponding cut-off ratio (0.11) maximized the accuracy in predicting amyloid-positivity as determined by Pittsburgh compound B (PiB) PET. Amyloid groups were further divided into clinical groups according to their CDR scores as shown in Table 3.

### Brain tissue homogenization

Frozen brain tissue from the Charles F. and Joanne Knight ADRC was sliced using a cryostat at -20, weighed, collected in tubes, and stored at -80 prior to biochemical analyses. Brain homogenization buffer (25 mM tris-hydrochloride pH 7.4, 150 mM sodium chloride, 10 mM EDTA [ethylenediaminetetraacetic acid], 10 mM EGTA [ethylene glycol-bis{β-aminoethyl ether}-N,N,N′,N′-tetraacetic acid], phosphatase inhibitor cocktail [Sigma], and protease inhibitor cocktail [Roche]) was added to each tissue sample at a concentration of 0.1 mg/µL of brain tissue at 4, and tissue was sonicated (Fisherbrand™ Model 120 Sonic Dismembrator) for 15 sec with 1 sec on and 1 sec off pulses using 35 % intensity. Once the samples were homogenized, they were centrifuged at 11,000 x g for 20 minutes at 4, to pellet cell/tissue debris. 100µL aliquots of supernatant were carefully transferred into new 0.6 mL Axygen tubes. The remaining supernatant and pellet were stored at -80 °C.

### Brain, spinal cord and peripheral nerve homogenization

Frozen brain, spinal cord and sciatic nerve samples from the ALS postmortem core and young control donors were cryo-sectioned, weighed, and collected in tubes. Cold brain homogenization buffer was added to the tissue samples at 3.25 μL buffer/ mg tissue and sonicated (Fisherbrand™ Model 120 Sonic Dismembrator) for 15 sec with 1 sec on and 1 sec off pulses using 35 % intensity. After sonification, samples were centrifuged at 11,000 x g for 20 minutes at 4 (Eppendorf Centrifuge 5424R) to pellet debris. Supernatant was carefully removed and aliquoted into new tubes. Sarkosyl (N-Lauroylsarcosine sodium salt, Sigma) was added to the supernatant to a final dilution of 1% sarkosyl and allowed to incubate for 60 min prior to ultracentrifugation at 100,000 g for 1 hour at 4 (Beckman Optima TLX). The supernatant was aliquoted into new tubes and stored at -80 prior to use.

### Immunoblotting

Soluble lysate samples (20 µg) were mixed with 4X LDS sample buffer (Bio-Rad) containing a final concentration of 10 mM of 1,4-dithiothreitol (DTT, Pierce^TM^, A39255) and heated at 70 °C for 10 minutes. Samples were loaded into each well and electrophoresed at 120 V for 90 min on 4–12% NuPAGE^TM^ Bis-Tris mini protein gel (Invitrogen^TM^, NP0322) and then transferred to 0.45 μm PVDF membrane (Immobilon®, IPVH00010) and blocked for 1 hour at RT in 5% nonfat milk in phosphate buffer saline with 0.1% Tween20 (PBS-T). All antibodies were diluted in PBS-T containing 5% nonfat milk. Membranes were probed with primary monoclonal anti-tau antibodies Tau-1 (provided by Dr. Nicholas Kanaan’s lab from Michigan State University; 1:1000 dilution), Tau-5 (Chemicon®, MAB361, 1:1000 dilution), and loading controls monoclonal GAPDH (Invitrogen^TM^, MA5-15738, 1:2000), anti β-actin (Cell signaling technology, 13E5, 1:2000) overnight at 4°C. The membranes were washed in PBS-T for 3 cycles (3 × 10 min) and incubated in goat anti-mouse IgG-HRP linked secondary antibody (Invitrogen^TM^, A28177, 1:2000), or goat anti-rabbit IgG-HRP linked secondary antibody (Cell signaling technology, 7074, 1:2000) for 1 hour at RT and washed with PBS-T. Immunoblot of cortical and cerebellar brain lysates were developed using West Pico PLUS chemiluminescent substrate (SuperSignal^TM^, Cat#34580) and imaged for 120 sec. In contrast, immunoblots of peripheral nerve lysates were developed using West Atto ultimate sensitivity substrate (SuperSignal^TM^, Cat#A38554) and imaged at high exposure (120 sec) to enhance visualization of low-abundance HMW and LMW tau species expressed in the periphery using Bio-Rad Chemidoc Imaging System.

### SDS-PAGE and in-gel protein digestion

Sarkosyl soluble homogenates (20 µg) from CNS and PNS were reconstituted with 4X LDS sample buffer (BioRad) containing 10 mM of reducing agent dithiothreitol (DTT). Each tissue sample was loaded into an individual well on a 4–12% NuPAGE^TM^ Bis-Tris mini protein gel. The gel was electrophoresed at 120 V for 90 min. After electrophoresis, the gel was stained with Oriole fluorescent gel stain (Bio-Rad, Cat#1610496) for 30 min in dark. Gel was imaged using Bio-Rad Chemidoc Imaging System. Gel image was printed out and regions to cut were noted and numbered. Each gel band was cut using a clean scalpel (Med PRIDE) for in-gel digestion using trypsin protease. The gel bands were de-stained using 25 mM ammonium bicarbonate (NH4HCO3) in 50% Acetonitrile (ACN) solution and completely dried, reduced with 10 mM DTT at 60°C for 30 min and alkylated in the dark using 20 mM iodoacetamide (IAA, Pierce^TM^, A39271) solution at RT for 60 min. Gel bands were dehydrated in the vacuum concentrator and rehydrated using a 40 µL of 10 ng/μL trypsin solution (25 mM TEABC pH 8.0) on ice for 10 min. The gel bands were digested for 18 hours at 37 °C and the supernatant was desalted on Oasis μElution HLB plates (Waters).

### Soluble tau quantitation in human tissues and CSF

Total protein concentrations of human tissue homogenates were determined using the bicinchoninic acid (BCA) assay method.^59^ For the soluble tau analysis, tau was immunoprecipitated (IP) using Tau-1, HJ8.5 and HJ8.7 antibody mixture as previously described with adjustments.^14,56^ To each soluble brain supernatant (50 µg total protein), 0.01% HSA solution was added along with 0.625 ng of ^15^N-2N4R (0.15 ng/µL, gift from Dr. Guy Lippens, France) and 1.25 ng of ^15^N-0N3R (0.15 ng/µL, Promise Proteomics, Grenoble, France) to a final volume of 500 µL. Tau was immunoprecipitated from 0.5 mL CSF. Soluble tau from each sample (brain or CSF) was immunoprecipitated in detergent (1 % NP-40), chaotropic reagent (5 mM guanidine), and protease inhibitors (Roche), using the antibody cocktail (50 % slurry of tau antibody conjugated sepharose (cyanogen bromide activated) beads containing 3 µg antibody/mg beads) by incubating them with rotation for 2 hours at room temperature. Following incubation, the samples were washed three times with 25 mM TEABC (Triethylammonium bicarbonate, Sigma-Aldrich, St. Louis) buffer. The immobilized proteins were digested on beads for 18 hours using 0.4 µg trypsin (Promega) at 37 . 50 fmol each of AQUA internal standard peptide for unmodified tau peptides, 5 fmol of phospho-tau AQUA internal standard and 25 fmol of tau exon 4a and exon 6 AQUA internal standard peptides were spiked in the digested samples for their quantification. Soluble tau digests were desalted using the Oasis µElution HLB plate (Waters) according to manufacturer’s protocol. The eluent was lyophilized and reconstituted with 25 µL of 2 % ACN, 0.1 % FA in water prior to MS analysis on Vanquish Neo UHPLC (Thermo Fisher Scientific, USA) coupled to Orbitrap Exploris 480 (Thermo Fisher Scientific, USA). Eighteen common tau peptides were quantified by comparison with the corresponding isotopomer signals from the ^15^N internal standard using Skyline software (version 20.2 MacCoss Lab, Department of Genome Sciences, University of Washington). The exon 4a-L and exon 6 peptides were quantified by comparison with the corresponding isotopomer signals from the AQUA peptides. Relative quantification for common tau peptides, exon 4a-L and exon 6 peptides were calculated by taking the “absolute” amount for each peptide and dividing by the “absolute” amount of the reference peptide TPSLPTPPTR (referred to in this manuscript as the “total-tau peptide”). The protein profile results are plotted as a function of peptide amino acid start position-end position and each point represents the mean for the given group, with error bars indicating the standard error of mean (SEM).

To validate the endogenous junction peptides for exon 4 to exon 4a-L (Table 2 and Supplementary Table 1) and exon 4 to exon 4a-S, custom synthesized AQUA peptides for 0N-4a-L (AEEAGIGDTPSLEDEAAGHVTQEELRVPGRQR), 1N-4a-L (STPTAEAEEAGIGDTPSLEDEAAGHVTQEELRVPGRQR), 2N-4a-L, (QAAAQPHTEIPEGTTAEEAGIGDTPSLEDEAAGHVTQEELRVPGRQR) and 0N-4a-S (AEEAGIGDTPSLEDEAAGHVTQEPESGKVVQEGFLR), 1N-4a-S, (STPTAEAEEAGIGDTPSLEDEAAGHVTQEPESGKVVQEGFLR) and 2N-4a-S (QAAAQPHTEIPEGTTAEEAGIGDTPSLEDEAAGHVTQEPESGKVVQEGFLR) were purchased from Life Science Technologies (Thermo Fischer Scientific). 50 ng of each of these AQUA peptides (concentrated stock 10 ng/µL prepared in 1 % ACN, 0.1 % FA) were spiked into 0.1% HSA solution along with 5 ng of ^15^N-2N4R and 5 ng of ^15^N-0N3R to a final volume of 500 µL. Tau immunoprecipitation followed by tryptic digestion and LC-MS was performed as described above. The LC-MS traces of the synthetic trypsin digested peptides (Supplementary Table 2) were compared with those of their respective endogenous tau junction peptides derived from human tissue lysates.

**Table 2.**
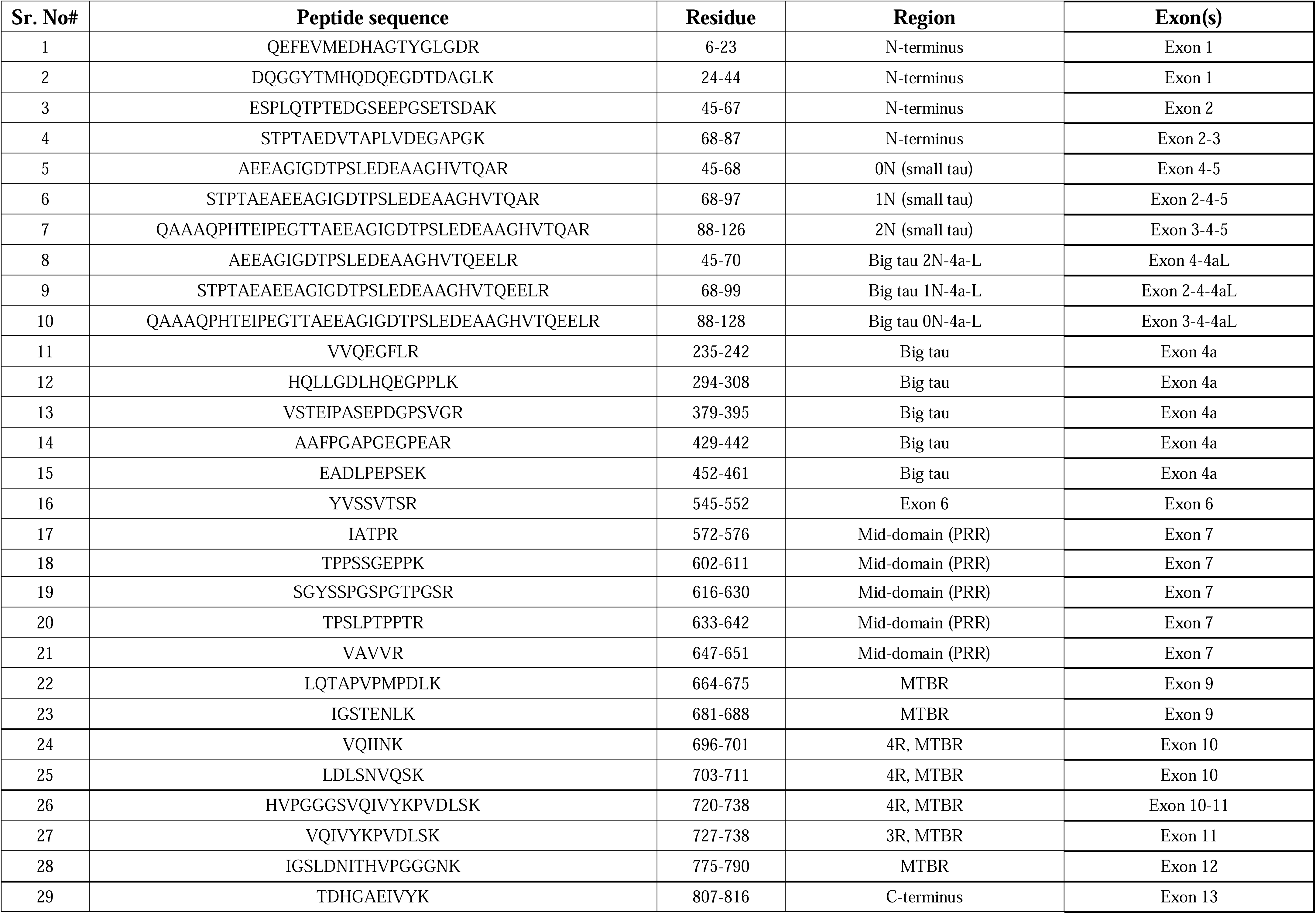
The list of exon 4a-L, big tau variant isoforms tryptic peptides quantified in this study by using human nervous tissues. Microtubule binding region, MTBR; proline rich region, PRR.

**Table 3.** Demographics and biomarker values of the human CSF study participants.

| Variable | Young Normal Controls | CDR = 0 |  | CDR > 0 |  |
| --- | --- | --- | --- | --- | --- |
| | | Aβ <sup>-a</sup> | Aβ <sup>+b</sup> | Aβ <sup>-c</sup> | Aβ <sup>+d</sup> |
| N | 25 | 25 | 14 | 4 | 21 |
| Gender, (F/M) | 13/11 (1 NA) | 13/11 (1 NA) | 5/9 | 3/1 | 10/11 |
| Age (years) | 42.45 ± 13.17 | 72.35 ± 4.68 | 73.14 ± 5.55 | 71.24 ± 1.75 | 76.92 ± 7.03 |
| CDR<sup>e</sup> | NA | 0 | 0 | 0.5 | 0.5-2 |
| MMSE<sup>f</sup> | NA | 29.25 ± 0.97 | 29.33 ± 0.78 | 28.75 ± 0.50 | 23.75 ± 5.29 |
| CSF Amyloid-β 42/40 | 0.13 ± 0.03 (1 NA) | 0.14 ± 0.02 (1 NA) | 0.09 ± 0.01 | 0.14 ± 0.01 | 0.07 ± 0.01 |
| pT181/T181 (%) | 6.89 ± 1.21 | 8.08 ± 2.23 | 8.13 ± 2.13 | 6.52 ± 2.90 | 9.61 ± 1.67 |
| pT217/T217 (%) | 1.55 ± 0.66 | 1.52 ± 0.53 | 3.11 ± 1.24 | 1.83 ± 0.36 | 6.37 ± 1.78 |
| MTBR-tau243 (ng/mL) | 0.12 ± 0.05 | 0.18 ± 0.07 | 0.22 ± 0.05 | 0.12 ± 0.04 | 0.55 ± 0.28 |
| T181 (ng/mL) | 4.22 ± 1.74 | 5.77 ± 2.27 | 6.49 ± 1.38 | 3.90 ± 0.77 | 9.33 ± 2.54 |
| T212 (ng/mL) | 3.20 ± 1.43 | 4.31 ± 1.81 | 4.96 ± 1.20 | 2.77 ± 0.53 | 6.72 ± 2.05 |
| Big tau-235 (fmol/mL) | 0.03 ± 0.03 | 0.07 ± 0.04 | 0.06 ± 0.03 | 0.06 ± 0.03 | 0.08 ± 0.06 |
| Big tau-294 (fmol/mL) | 0.06 ± 0.06 | 0.10 ± 0.06 | 0.08 ± 0.06 | 0.09 ± 0.10 | 0.12 ± 0.09 |
| Big tau-429 (fmol/mL) | 0.02 ± 0.02 | 0.04 ± 0.02 | 0.04 ± 0.02 | 0.05 ± 0.03 | 0.06 ± 0.04 |
| Big tau-452 (fmol/mL) | 0.02 ± 0.02 | 0.04 ± 0.02 | 0.04 ± 0.02 | 0.04 ± 0.02 | 0.05 ± 0.03 |
| Big tau-545 (fmol/mL) | 0.004 ± 0.005 | 0.006 ± 0.004 | 0.008 ± 0.007 | 0.003 ± 0.002 | 0.015 ± 0.011 |
| Small tau45-67 (0N), (ng/mL) | 1.62 ± 0.83 | 2.05 ± 0.92 | 2.51 ± 0.66 | 1.45 ± 0.56 | 3.41 ± 1.03 |
| N-term, tau6-23 (ng/mL) | 1.20 ± 0.57 | 1.65 ± 0.71 | 2.11 ± 0.40 | 1.05 ± 0.24 | 3.44 ± 1.02 |

Data presented as mean ± standard deviations (SD); <sup>a</sup> Age-matched controls; <sup>b</sup> Preclinical-AD non-AD cognitively impaired; <sup>d</sup> mild to moderate-AD; <sup>e</sup> Clinical Dementia Rating scores Mini Mental State Examinations; NA Data not available

CSF tau MTBR containing the residue 243 (MTBR-tau243) peptides were monitored using the previously reported procedures with minor modifications.^15^ Briefly, 0.45 mL of post-IP (for big tau species described in the previous section) CSF was thawed and 5 μL of ^13^C^15^N 2N4R-tau internal standard (250 pg/µL) spiked into the post-IP CSF. Then, the tau species containing the residue 243 were immunoprecipitated with HJ32.11 antibody.^14^ The immunoprecipitated tau species were digested by trypsin and desaltated by the Oasis µElution HLB plate (Waters) according to manufacturer’s protocol. The samples were reconstituted with 25 µL of 3 % ACN, 3 % FA in water and peptides were separated by nanoAcquity ultra-performance LC system (Waters) which was coupled to Orbitrap Tribrid Eclipse (Thermo Fischer Scientific, USA) mass spectrometer operating in parallel reaction monitoring mode (PRM). The resulting MTBR-tau243 peptide (residue 243-254) and the corresponding isotopomer from the ^13^C^15^N internal standard were monitored and quantified similarly as other tau species including big-tau as described above.

### Mass Spectrometry

The peptides were directly loaded on to HSS T3 75 μm × 100 mm, 1.8 μm C18 column (Waters) heated to 65 using a Vanquish Neo UHPLC (Thermo Fisher Scientific, San Jose, CA, USA) and peptides were separated using a flow rate of 0.4 µL/min with a mixture of buffer A (0.1 % FA in water) and buffer B (0.1 % FA in acetonitrile). Tau peptides were eluted from the column with a gradient of 4 %-8 % of buffer B for 10 min, then 8 %-20 % of buffer B for another 8 min before ramping up to 95 % buffer B in next 2 min and cleaning the column for another 2 min. The Thermo Orbitrap Exploris 480 was equipped with a Nanospray Flex electrospray ion source (Thermo Fisher Scientific, San Jose, CA, USA) and operated in positive ion mode. Peptide ions sprayed from a 10 μm SilicaTip emitter (New Objective, Woburn, MA, USA) into the ion source (spray voltage = 2.2 kV), were targeted and isolated in the quadrupole. Isolated ions were fragmented by high-energy collisional dissociation (HCD), and ion fragments were detected in the Orbitrap (resolution of 15,000 for tau, 30,000 for phospho-tau and 120,000 for big tau peptides, mass range 150–1,500 m/z).

### LC-MS/MS for bottom-up proteomics

Peptides were loaded directly using HSS T3 75 μm × 100 mm, 1.8 μm C18 column (Waters) heated to 65 using a Vanquish Neo UHPLC (Thermo Fisher Scientific, San Jose, CA, USA) and peptides were separated using a flow rate of 0.4 µL/min with a mixture of buffer A (0.1 % FA in water) and buffer B (0.1 % FA in acetonitrile). Peptides were eluted from the column with a stepped gradient of 52 min starting from 0.5 % buffer B to 15 % buffer B, 35 min; 30 % buffer B, 52.5 min. The column was washed with 80 % buffer B for 3 min before equilibration at 0.5 % buffer B for 5 min. The proteolytic peptides were sprayed into MS inlet at 2.2 kV in positive ion mode using commercial PicoTip emitters (New Objective). A full mass spectrum scan with a resolution of 120,000 @ *m/z* 400 was acquired in the mass range *m/z* 350-1500 (AGC target 2×10, maximum injection time 54 ms). Peptides were selected for fragmentation using higher-energy collisional dissociation (HCD) using MIPS (monoisotopic precursor selection) mode. MS2 scan settings were the following: resolution 22,500, AGC target 5 ×10^5^, maximum injection time 22 ms, isolation window 1.2 *m/z*, normalized collision energy 30 %, filter intensity (intensity threshold 5e^4^), charge states 2-5 selected, undetermined charge states filter on, exclude isotopes on, and dynamic exclusion 30 s.

Raw data was analyzed in Fragpipe using human proteome database downloaded in January 2024 from UniProt (only reviewed entries and appended with in-house fasta containing manually curated tau exon-4a-L and exon 4a-S sequences). Currently only exon 4a-S containing big tau sequence is present in UniProt – P10636-1. We created the in-house fasta along with P10636-1 (tau with exon 4a-S, NCBI Ref seq ID: NP_058519) along with the sequences curated from NCBI protein database (XP_005257423, XP_001364194, XP_005257419) corresponding to three isoforms of tau containing exon 4a-L. The raw data were searched with this curated protein database using MSFragger database search engine.^60^ The following settings were applied: trypsin (specificity set as C-terminal to arginine and lysine) with up to three missed cleavages, the mass tolerance was set to ± 20 ppm for the precursor and fragment mass tolerance. Fixed modifications included Cys carbamidomethylation (CAM, mass shift +57.0215 Da) with variable modifications including Ser / Thr / Tyr phosphorylation (+79.9663 Da), Gln and Asn deamidation (+0.98 Da) and Met oxidation (+15.9949 Da). False discovery rate (FDR) was set to 1% on peptide and protein levels (PeptideProphet and Philosopher)^61^ with a minimum length of five amino acids and was determined by searching a decoy reverse database. For all other search parameters, default settings were used. Similar parameters were used for the trypsin + Asp-N and tryspin + Glu-C digested samples with the setting for the corresponding dual enzyme setting. Label-free quantification was done using the in-built label-free quantification (IonQunat) algorithm integrated in MSFragger search.^62^

### *MAPT* Targeted IsoSeq

Long read RNA sequencing on *MAPT* (IsoSeq) was performed as previously described.^63^ Briefly, we used a modified single molecule real time bell-shaped template (SMRTbell) amplicon protocol at the Center for Advanced Genomic Technologies, Mount Sinai. cDNA synthesis was performed from 100 ng RNA per sample with SuperScript IV First-Strand Synthesis System (ThermoFisher Scientific, Cat. No.18091200) with oligo(dT) primers. MAPT-specific PCR amplification employed TaKaRa LA Taq DNA Polymerase with GC Buffer (Clontech, Cat. No. RR02AG) and gene-specific primers MAPT_NF_F1 (5′–ATG GAA GAT CAC GCT GGG AC–3′) and MAPT_NF_R2 (5′–GAG GCA GAC ACC TCG TCA G–3′). Amplicons were purified using AMPure PB beads (PacBio) followend-repair, A-tailing and ligation of barcoded SMRTbell adapters using the SMRTbell Express Template Prep Kit 3.0 (PacBio). After cleanup, final libraries were sequenced on the PacBio Sequel II platform using circular consensus sequencing (CCS) to generate high-fidelity (HiFi) reads. Demultiplexing and adapter trimming were performed with lima, and artificial concatemers were removed. Isoform clustering and polishing were done using the Iso-Seq tool suite, and alignments were performed against GL000258.2 and KI270908.1 references using pbmm2 with the ISOSEQ preset. Redundant isoforms were collapsed and classified using pigeon, referencing GENCODE v39 annotations. Low-confidence and single-exon transcripts were excluded. The resulting isoforms were converted to BED format for visualization and cross-referenced against an internal isoform atlas to maintain consistent naming. Lastly, a computational pipeline was used to process the BED files, naming transcripts based on presence of known exons, and plot them to evaluate 4a/4aL expression and isoform composition across brain regions.

### Data and Statistical Analysis

Data are represented as mean ± standard deviation (SD) unless mentioned otherwise and were plotted in GraphPad Prism (version 10.5). One-way ANOVA followed by post analyses (Tukey’s or Dunnett’s t-tests as appropriate) were performed to compare between multiple brain regions and CSF tau levels. Significance was evaluated at the 0.01 and 0.05 level. Spearman correlations were used to assess the associations between CSF tau biomarkers, age, and exon 4a-L and exon 6 peptides. Non-parametric tests were used for non-normally distributed data. Spearman correlation was used for continuous variables. Diagnostic performances were evaluated with receiver operating curves (ROC) and area under the curve (AUC) assessments.

## Results

### Tau exon 4a-long translation leads to insertion of 355 amino acids in human “big tau”

Tau transcript could be extended by the insertion of exon 4a (between exons 4 and 5) and exon 6 (Figure 1). Big tau is distinguished from canonical small tau isoforms based on the inclusion and translation of exon 4a and exon 6. The tau isoform repertoire beyond the CNS is further diversified by the inclusion of only exon 4a in the tau transcript which has been described to be as exclusively abundant in the periphery – “PNS tau” (Figure 1A). Alternative 3’ splicing of tau exon 4a leads to a larger splice variant (exon 4a-L) in human cancer cells and skeletal muscle tissue (Figure 1b).^45,46^ However, two different splice variants of exon 4a have been annotated, UniProt ID:P10636-1 for exon 4a, referred to as exon 4a-short (exon 4a-S) and NCBI ID: XP_005257419 for exon 4a-long (exon 4a-L) (Figure 1a-c).^50^ These splice variants could theoretically lead to insertion of either 251 (exon 4a-S) or 355 (exon 4a-L) amino acids in the N-terminal projection domain of tau (Figure 1a, 1c) in humans, respectively.^46^ Thus, to determine which big tau protein isoforms are expressed in adult human nervous tissue (Supplementary Figure 1a), we defined their amino acid sequences using mass spectrometry (MS).

**Figure 1:**
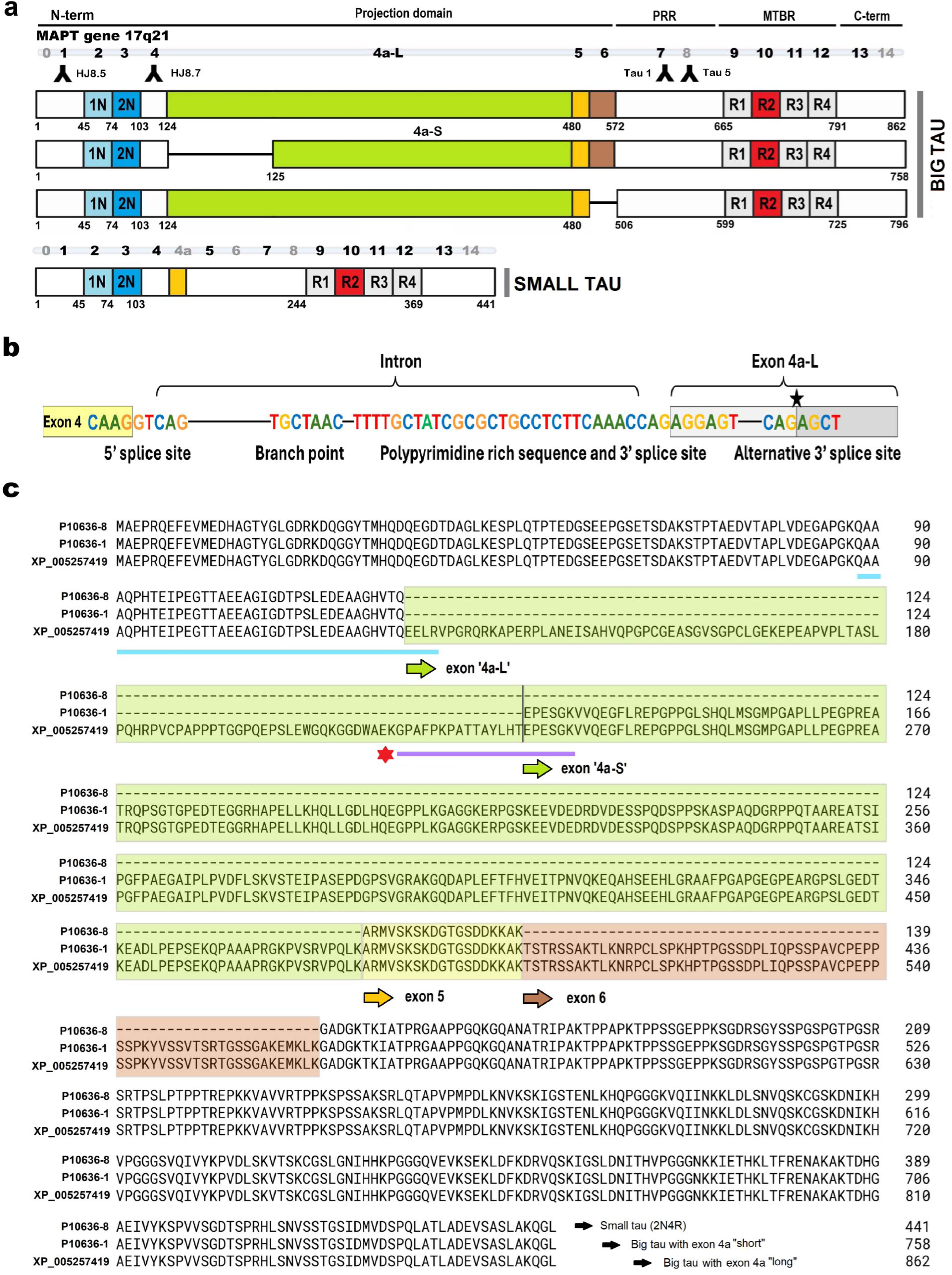
Big tau - inclusion of the exon 4a-long and exon 6 leads to the longest *MAPT* isoform sequence. **a.** Schematic representation of the full-length human tau isoforms with and without exon 4a. Insertion of exon 4a-L and exon 6 result in “big tau” that is 862 amino acids long (top). The insertion of exon 4a-S and exon 6 will result in big tau isoform with a 758 amino acid sequence (second from top). Big tau isoform can also result from the insertion of exon 4a-L without exon 6 (third from top). Canonical small tau (2N4R) lacks both the exon 4a and exon 6. Monoclonal anti-tau antibodies targeting, N-terminal and proline rich region (HJ8.5, HJ8.7, Tau-1, Tau-5) are depicted on the sequence. **b.** Human *MAPT* genomic sequence highlighting 5′ splice site of the exon 4, branch point sequence (BPS), polypyrimidine rich sequence, canonical 3′ splice site, and alternative 3′ splice site within exon 4a-L (black star), indicating regulatory elements that promote exon 4a-L insert. **c.** The amino acid sequence of the longest human big tau proteins with either exon 4a-L or exon 4a-S aligned with the longest common tau isoform found in the adult human brain (2N4R). The N-terminus of exon 4a-L is highlighted by green arrow, while the N-terminus of the exon 4a-S is depicted by the vertical black line on the amino acid sequence. Tryptic peptide for exon 4 to exon 4a-L junction peptide (2N-4a-L peptide) is highlighted in the second line. Asterix highlights the exon 4a-L tryptic peptide (big tau 4a-L*) corresponding to alternative 3’ splicing.

Using a combination of antibodies targeting tau N-terminus (HJ8.5 and HJ8.7) and mid-domain (Tau-1) specific epitopes, we immunoprecipitated tau from the tissue lysates originating from CNS and PNS. To determine which big tau isoforms are expressed in adult human nervous tissue (Supplementary Figure 1a), we defined their amino acid sequences using mass spectrometry (MS). We performed bottom-up tandem mass spectrometry (MS/MS) on pooled homogenates from adult human brain, spinal cord, and sciatic nerve, respectively. Proteins were digested into peptides and analyzed by liquid chromatography–tandem mass spectrometry (LC–MS/MS). Peptide spectra were searched against a customized protein database containing tau sequences incorporating both exon 4a-short (4a-S) and exon 4a-long (4a-L) variants. This strategy enabled the identification of exon 4a–specific peptides and exon-bridging peptides unique to respective big tau isoforms. Detection of these peptides provided direct protein-level evidence for the translation of exon 4a-containing tau isoforms in human nervous tissue and allowed us to distinguish between the predicted 4a-L and 4a-S splice products (Figure 2a). We observed the presence of tryptic peptides specific to the translation of exon 4a in the brain, spinal cord and sciatic nerve, respectively (Supplementary Figure 1b). Multi-enzyme digestion (Supplementary Figure 1b-1c) further validated exon 4a expression at the protein level. The insertion of both the exons 4a-L and 6 would result in an 862 amino acid long protein sequence of tau, making it the longest isoform of human tau — “big tau” (Figure 1c). The higher-energy collision dissociation (HCD)-MS/MS spectrum of the tryptic big tau peptide derived from the union of exon 4 and exon 4a-L (big tau 2N-4a-L) (Figure 2b), [M+4H]^4+^ *m/z* 1064.7520 with the *b* (*b*_2_-*b*_20_) and *y* ion series (*y*_1_-*y*_26_) validates the peptide amino acid sequence “QAAAQPHTEIPEGTTAEEAGIGDTPSLEDEAAGHVTQEELR” (Figure 2b). This tryptic peptide is different from the sequence “QAAAQPHTEIPEGTTAEEAGIGDTPSLEDEAAGHVTQAR” found in the common small tau isoforms (exon 4-exon 5 junction) without exon 4a-L insert (Figure 2a, Supplementary Figure 2A-C). We found another peptide with “GPAFPKPATTAYLHTEPESGK” (Figure 2c) sequence that is specific to exon 4a-L. The human exon 4a-L has been described as the exon 4a-S plus additional coding sequence at the 5’ end of exon 4a-S (Figure 2a).^45,46^ Identification of the “GPAFPKPATTAYLHTEPESGK” peptide (Figure 2c) provides crucial evidence of the alternative 3’ splice junction within the exon 4a-L.^39^

**Figure 2.**
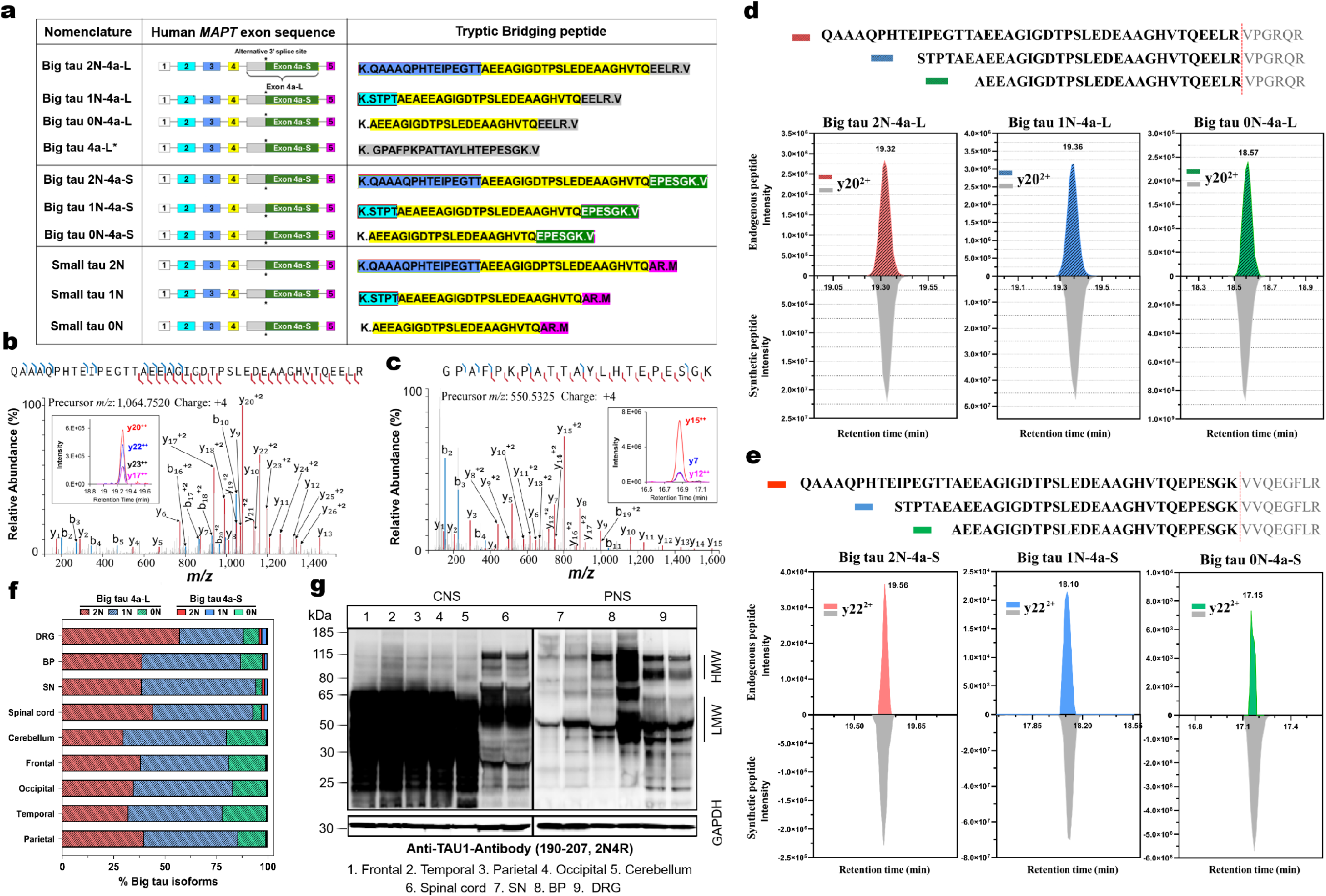
Exon bridging peptides for identification of big tau isoforms in humans. **a.** Schematic representation of the *MAPT* exon structure (exons 1–5) illustrating isoforms with and without exon 4a, together with the corresponding amino acid sequences of tryptic bridging peptides. Tryptic bridging peptides from 2N, 1N, and 0N big tau isoforms generated by insertion of exon 4a-L are distinct from those arising from exon 4a-S insertion. For reference, the canonical small tau 2N, 1N, and 0N peptides are derived from exon 4–exon 5 junctions. **b.** Representative HCD-MS/MS spectrum of the unique big tau 2N-4a-L bridging tryptic peptide [M+4H]^4+^ *m/z* 1064.7520. The inset shows the extracted ion chromatograms (XIC) depicting the distribution of top 4 fragment ions from the MS/MS spectra. **c.** Representative HCD-MS/MS spectrum of big tau 4a-L* specific tryptic peptide [M+4H]^4+^ *m/z* 550.5325 corresponding to alternative 3’ splicing (right). The *y* and *b* ions are shown in red and blue, respectively. The inset shows the extracted ion chromatograms (XIC) depicting the distribution of top 3 fragment ions from the MS/MS spectra. **d.** Representative mirror plots for XIC for three endogenous big tau exon 4a-L bridging peptides (top) following trypsin digestion showing complete match with their respective synthetic standard peptides (bottom) on the LC-MS. The *y*_20_^2+^ ion (most abundant fragment ion) was used for comparison between respective endogenous and standard peptides. The peptide sequence for the three exon 4a-L isoforms (0N-, 1N-, 2N-4a-L) that get generated following trypsin digestion are depicted above the XIC plots. **e.** Representative mirror plots for XIC for the endogenous big tau exon 4a-S bridging peptides following trypsin digestion showing complete match with their respective synthetic standard peptides (bottom) for 2N, 1N and 0N isoforms on the LC-MS. The *y*_22_^2+^ ion (most abundant fragment ion) was used for comparison between respective endogenous and standard peptides. The peptide sequence for the three exon 4a-S isoforms (0N-, 1N-, 2N-4a-S) that get generated following trypsin digestion are depicted above the XIC plots. Brachial plexus tissue was used for (c) and (d) for generating these mirror plots. **f.** Relative distributions (%) of respective big tau isofroms (exon 4a-L+exon 4a-S) quantified in CNS (parietal, temporal, occipital, frontal, cerebellum, spinal cord, each n=6), and PNS (DRG, n=2; BP, n=2 and SN, n=5) tissues. Abbreviations: DRG, dorsal root ganglion; BP, brachial plexus; SN, sciatic nerve. **g.** Representative immunoblot analysis of tau protein isoforms across CNS and PNS tissues in humans, using Tau-1 antibody. Big tau (HMW tau ∼ 115-90 kDa) and small tau (LMW tau ∼ 65-40 kDa) protein expression were detected in the PNS (SN, DRG and BP) and spinal cord tissues. No discernable HMW tau bands were detected across brain regions (parietal, temporal, occipital, frontal and cerebellum). GAPDH (35 kDa) was used as loading control. Abbreviations: DRG, dorsal root ganglion; BP, brachial plexus; SN, sciatic nerve; GAPDH, glyceraldehyde 3-phosphate dehydrogenase.

In canonical small tau, the alternative splicing of exons 2 and 3 with exon 4 predominantly generates 0N and 1N isoforms, with less abundant 2N isoform found in the brain (Supplementary Figure 2A-C).^27,64^ This led us to investigate whether alternative splicing of exons 2 and 3, together with the expression of exon 4a-L or exon 4a-S, could generate additional isoforms akin to small tau isoforms (Figure 2a). Indeed, along with the big tau 2N-4a-L (Figure 2b) peptide we identified specific peptides corresponding to 0N-4a-L, 1N-4a-L isoforms (Supplementary Figure 2D-E, 2G-I). Synthetic standard peptides (Supplementary Table 2) for the corresponding bridging peptides for big tau isoforms (4a-L and exon 4a-S) were used for orthogonal validation (Supplementary Figure 3). Tau immunoprecipitation followed by tryptic digestion of the synthetic big tau 2N-4a-L, 1N-4a-L and 0N-4a-L peptides matched the corresponding endogenous big tau peptides by LC-MS (Figure 2d), confirming the identity of big tau isoforms in human samples. Comparison of the LC-MS analysis of the synthetic big tau 4a-S peptides confirmed that the corresponding 2N-4a-S, 1N-4a-S and 0N-4a-S endogenous peptides (Figure 2e, Supplementary Figure 3) can be also detected in human nervous tissues, albeit at low levels. We observed across human CNS (parietal, temporal, frontal, occipital, cerebellum and spinal cord) and PNS (sciatic nerve, brachial plexus, and dorsal root ganglion), the relative abundance of big tau 4a-L isoforms (0N-4aL+1N-4a-L+2N-4a-L) greatly exceeded that of 4a-S isoforms (0N-4a-S+1N-4a-S) (Figure 2f and Supplementary Figure 3B). We estimated big tau exon 4a-L isoforms (> 99% in CNS brain regions and > 97% in CNS, spinal cord and PNS, sciatic nerve, brachial plexus, > 96.5 % in PNS, dorsal root ganglion) >> big tau exon 4a-S isoforms (< 1 % in CNS brain regions and < 3.5 % in CNS, spinal cord and PNS regions, sciatic nerve, brachial plexus, dorsal root ganglion) (Figure 2f). Together, these results demonstrate that big tau in the human nervous system is generated predominantly through insertion of exon 4a-L (> 95%) between exons 4 and 5, whereas exon 4a-S insertion (< 5%) provides a minor contribution to big tau isoforms. (Figure 2f and Supplementary Figure 3).^50,55^ Our observation aligns with what has been previously reported in human cancer lines and skeletal muscles tissues.^46^ Along with exon 4a-L (in the rest of the paper, we have focused on these big tau isoforms) expression across the nervous tissue, we also observed exon 6 peptides across CNS and PNS tissue lysates (Supplementary Figure 2C and Supplementary Figure 2F).

Big tau protein, twice as large compared to small tau, has been predicted to have molecular weight ∼ 110 kDa.^55,65,66^ Immunoblotting with tau antibodies targeting the proline rich region (Tau-1 and Tau-5, Figure 2g and Supplementary Figure 4) indicated the existence of high molecular weight (HMW tau, ∼ 115-90 kDa) tau bands, consistent with big tau, along with the common low molecular weight (LMW tau, ∼ 40-65 kDa) tau species. These HMW bands were detected in the human PNS tissues (sciatic nerve, brachial plexus and dorsal root ganglion) and selected CNS region (spinal cord tissues). Notably, no clear discernable HMW bands (∼ 115-90 kDa) were observed in select brain tissues (frontal, temporal, parietal, occipital and cerebellum) using immunoblotting (Figure 2g and Supplementary Figure 4). To determine whether the HMW tau species detected by immunoblotting contain exon 4a-L, we performed SDS–PAGE followed by in-gel digestion and LC–MS on nervous tissue lysates from the CNS and PNS (Supplementary Figure 5). Peptides corresponding to big tau exon 4a-L (2N-4a-L, 1N-4a- L and 0N-4a-L) were detected in the HMW (75-185 kDa) gel bands from both the CNS and PNS samples. Big tau exon 4a-L peptides were abundant in the HMW bands in the spinal cord, brachial plexus and dorsal root ganglion (Supplementary Figure 5). Relatively lower levels of big tau exon 4a-L isoforms were also observed in the brain HMW gel band in the in-gel digestion LC-MS (Supplementary Figure 5C). We also detected 3R and 4R specific peptides in the HMW bands from the CNS and PNS samples, demonstrating that big tau contains MTBR along with exon 4a-L (Supplementary Figure 5).

To complement the protein-level observations and determine whether big tau transcripts are expressed in the adult human brain, we performed long-read IsoSeq^63^ sequencing on caudate (n=5) and frontal cortex (middle frontal gyrus, n=5) from ten different control brain donors (Supplementary Table 3). Exon 4a-L transcripts were detected at low levels (Supplementary Figure 6), while exon 4a-S transcripts were not detected. The exon 4a-L transcripts detected in these two brains regions were 1N-4a-L-3R, 1N-4a-L-4R and 0N-4a-L-3R (Supplementary Figure 6A). Similar to the proteomic results, big tau transcripts comprised ∼ 0.5-1 % of total tau detected in the two brains regions investigated (Supplementary Figure 6B). Together,our discovery proteomics, in-gel digestion with LC-MS, and IsoSeq findings from select brain, spinal cord and PNS tissues highlight distinct distribution pattern of HMW tau isoform in the human CNS and PNS.

### Expression pattern of tau exon 4a-L and 6 peptides across CNS and PNS

Building on our discovery proteomics findings, we developed a targeted immunoprecipitation-parallel reaction monitoring (IP-PRM) mass spectrometry assay to quantify the distribution and stoichiometry of big tau isoforms across the human nervous system. To quantitatively measure big tau peptides alongside other tau peptides, we refined and applied the IP strategy with parallel reaction monitoring (PRM)-MS technique.^56,57^ Using a combination of antibodies targeting tau N-terminus (HJ8.5 and HJ8.7) and mid-domain (Tau 1) epitopes (Figure 3a) shared by all tau isoforms (used in the discovery experiments), we immunoprecipitated endogenous tau and quantitatively measured the profiles of tau isoforms using the IP-PRM assay.^36^ We quantified more than 29 peptides across the tau protein sequence, which included peptides that are common to tau isoforms (18 peptides), 3 peptides specific to isoforms lacking exon 4a-L translation (small tau isoforms with N-terminal inserts: 0N, 1N and 2N), 3 big tau peptides that are specific to exon4-exon4a-L (0N-, 1N- and 2N-4a-L big tau isoforms), five peptides within the exon 4a-L itself and one peptide from exon 6 (Table 2).

**Figure 3:**
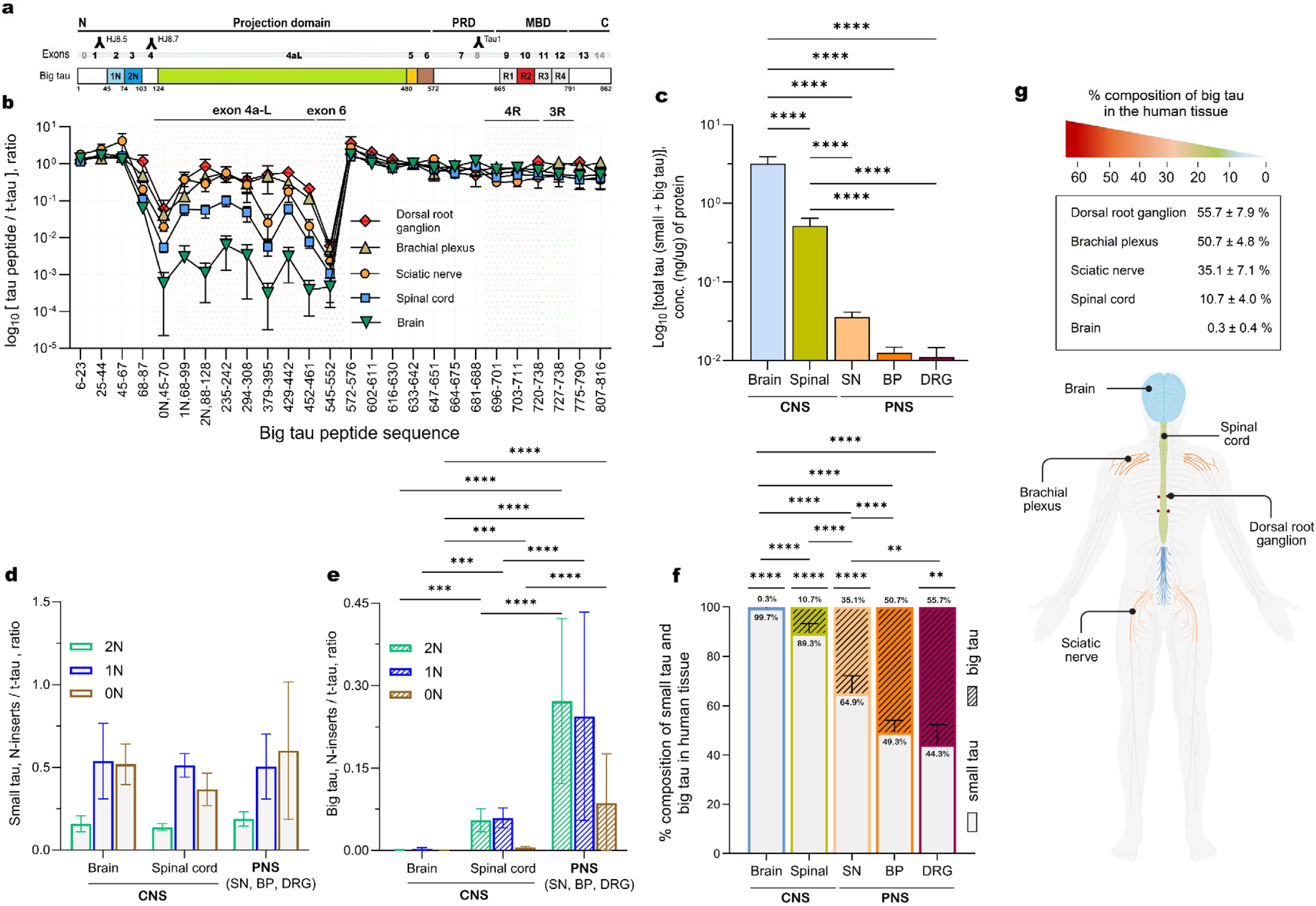
Differential expression of big tau across the human nervous system. **a.** Schematic of the big tau protein sequence indicating the N-terminal insert, proline rich domains and microtubule binding repeats. **b.** Quantitation of relative tau peptides including the exons 4a-L and 6 peptides from peripheral tissues, dorsal root ganglion (DRG, n=2, diamond), brachial plexus (BP, n=2, triangle) and sciatic nerve (SN, n=6, circle). Central nervous system, spinal cord (n=15, square) and brain (n=39, inverted triangle) tissue lysates. The tau peptides were normalized to total tau (residue 633-642 in reference to 2N big tau isoform). Big tau (exon 4a-L, 6), 4R and 3R specific peptides are highlighted. Data are presented as mean ± SD. **c.** Bar graphs represent the total tau levels (ng/µg protein) from all tau isoforms (big tau, PNS tau, small tau) were quantified in brain, spinal cord, sciatic nerve, brachial plexus and dorsal root ganglion samples used in the cohort in (**b**). Data was represented as mean ± SD. Statistical significance was performed using Brown-Forsythe and Welch ANOVA with post hoc multiple comparisons. ****p < 0.0001. **d.** Bar graphs of the three small tau isoforms (0N, 1N and 2N) normalized to t-tau for all the samples used in (**b**). Data are presented as mean ± SD. Statistical analysis was performed using two-way ANOVA with multiple comparisons, *p ≤ 0.05. **e.** Bar graphs of three big tau isoforms (0N, 1N and 2N big tau) normalized to t-tau for all the samples used in (**b**). Data are presented as mean ± SD, statistical analysis was performed using two-way ANOVA followed by multiple comparisons, ****p < 0.0001. **f.** Bar graph depicting the distribution of small tau and big tau N-terminal isoforms (0N+1N+2N) across the nervous system for all the samples used in (**b**). Data are expressed as mean ± SD. Statistical analysis was performed using two-way ANOVA with Tukey’s multiple comparisons test, ****p < 0.0001. **g.** Schematic diagram illustrating the relative big tau expression pattern across the human nervous system.

Typically, the shorter isoform of tau (small tau) has been referred as being CNS or brain specific while big tau (exon 4a-L) is exclusively found in the PNS and periphery.^16^ To test the hypothesis that exons 4a-L and 6 are differentially expressed across the nervous system, with higher expression in the PNS compared to the CNS,^39,40,47^ we quantified the common tau isoform peptides and big tau peptides in soluble tau extracted from six brain regions including parietal, temporal, occipital, frontal and cerebellum (CNS), spinal cord (CNS), and sciatic nerve, brachial plexus and dorsal root ganglion (PNS) samples (Figure 3b and Supplementary Figure 5, Table 1a and 1b). The quantitation of soluble tau peptides from the brain exhibited a profile expected for small tau isoforms as reported previously.^57^ We observed the exon 4a-L peptides in the brains, normalized to the mid-domain total-tau peptide chosen as reference (residue 633-642 according to big tau sequence (Figure 1c, 3b) or residue 212-221 according to small tau sequence (Figure 1d), were 200-fold lower than other common small tau peptides (Figure 3b, inverted triangle). The levels of exon 4a-L peptides (residue 235-242, 0.006 ± 0.005; residue 294-308, 0.003 ± 0.003; residue 429-442, 0.003 ± 0.002 compared to total-tau 633-642, n = 39) are two-orders of magnitude lower compared to other common tau peptides (residue 6-23, 1.379 ± 0.247; residue 45-67, 1.298 ± 0.362, residue 602-611, 1.019 ± 0.101 compared to total-tau) in the human brains (Figure 3b, inverted triangle and Supplementary Figure 5). These results indicate that the longest isoform of tau, “big tau” with exon 4a-L in the brain (globally combining all the brain regions studied) is present at lower proportion (∼ 0.3 %) compared to small tau (Figure 3b and 3f,). The exon 6 peptide (residue 545-552, 0.0005 ± 0.0033 compared to total-tau, n=39) indicates an even lower contribution to the total tau pool in the brain (less than 0.05 % of the total tau) (Figure 3b).

Compared to the brain extracts, we observed 20-fold more exon 4a-L peptide in the spinal cord lysates (Figure 3b, square). Amongst big tau peptides, the exon 4a-L peptide (residue 235-242) was the most abundant peptide in the spinal cord (0.100 ± 0.032 compared to total-tau, n = 15), an order-of-magnitude higher than what we detected in the brain lysates (*p* = 0.026) (Figure 3b). We also observed other exon 4a-L peptides (residue 294-308, 0.047 ± 0.021 and residue 429-442, 0.060 ± 0.017 compared to total-tau, n=15) were 20-fold higher in the spinal cord tissue (Figure 3b, square) samples compared to brain lysates. These results indicate that exon 4a-L is more widely expressed in spinal cord (∼ 10.7 % of total tau) compared to the brain (Figure 3b-3f). Compared to brain tissues, we detected 2.5-fold more exon 6 peptide (residue 545-552, 0.001 ± 0.001 compared to total-tau peptide, n=15) in spinal cord lysates (Figure 3b, square). These results highlight 10-fold higher exon4a-L expression compared to exon 6 in the spinal cord.

To further characterize the distribution of the tau exons 4a-L and 6 in the PNS, we measured soluble tau peptides from adult human sciatic nerve (SN, n = 6), brachial plexus (BP, n=2) and dorsal root ganglion (DRG, n=2) lysates. We detected exon 4a-L peptides at sub-equimolar amounts in the SN (residue 235-242, 0.566 ± 0.260; residue 294-308, 0.268 ± 0.180 and residue 429-442, 0.173 ± 0.082,n=6) (Figure 3b, circle), when normalized to total-tau (residue 633-642). We detected exon 4a-L peptides at similar ratios in BP (residue 235-242, 0.561 ± 0.250; residue 294-308, 0.294 ± 0.126 and residue 429-442, 0.295 ± 0.091 normalized to total-tau) and DRG (residue 0.441 ± 0.059; residue 294-308, 0.357 ± 0.202 and residue 429-442, 0.577-0.071 normalized to total-tau). This distribution profile indicates that big tau in the PNS constitutively expresses exon 4a-L. Our results highlight exon 4a-L is widely expressed in the PNS (∼ 50 %), 2-orders of magnitude higher compared to brain. We estimated low abundance of exon 6 peptide (residue 545-552) in the SN (0.002 ± 0.002), BP (0.005 ± 0.003) and DRG (0.006 ± 0.003) lysates (Figure 3b and Supplementary Figure 1c), reflecting a 10-fold increase compared to relative abundance in brain. Overall, these findings suggest there is a distribution pattern for tau exon 4a-L expression from the CNS to the PNS, increasing from very low levels (0.3 % of total tau) in the brain to intermediate levels in the spinal cord (10.7 % of total tau) to high levels (∼50 %) in the PNS.

Additionally, while the relative contribution of exon 4a-L to total tau increases from the CNS to PNS, we found that the concentration of total tau demonstrated an inverse gradient (Figure 3c). We estimated that brain contains ∼ 12 ng/µg protein of soluble tau (small and big tau combined), while there is 6-fold decrease (*p* < 0.0001) in the total tau levels in the spinal cord (∼ 2 ng/µg protein). The PNS (SN, BP and DRG) contains roughly 3 orders of magnitude lower levels of soluble tau (0.01-0.03 ng/µg protein) compared to brain (*p* < 0.0001). Additionally, we found exon 6 is expressed at a very low level globally compared to other common tau exons in the nervous system. The exon 6 expression is increased by 2.5-fold in the spinal cord (0.1 % of total tau) compared to the brain (less than 0.05 % of total tau), however we detected no significant changes in the sciatic nerve, dorsal root ganglion and brachial plexus (0.2 -0.5 % of total tau) compared to spinal cord (Figure 3b).

Along with the relative expression pattern of exon 4a-L across the nervous system, we also quantified the 3R and 4R isoform expression pattern as they are developmentally regulated.^27,67^ We observed a significant decrease in the 4R isoform (residue 720-738) levels in the spinal cord (0.49 ± 0.09 normalized to total-tau, *p* < 0.0001) and sciatic nerve lysates (0.38 ± 0.20, *p* < 0.0001) compared to brain (0.66 ± 0.23) (Figure 3b). However, 3R isoform (residue 727-738) remained unaltered in the spinal cord (0.47 ± 0.09) and sciatic nerve lysates (0.49 ± 0.23) compared to the brain (0.52 ± 0.20) (Figure 3b). These results further highlight that alternative splicing and insertion of tau exon 10 is also differentially regulated across the human nervous system.

### N-terminal projection of small and big tau across the nervous system

Adult human brain is known to express approximately 40 % 0N, 50 % 1N and 10 % 2N small tau isoforms (tau without exon 4a-L insert).^27,68,69^ However, the distributions of “big tau” isoforms across the nervous system have not been investigated. Guided by the IP-MS/MS data (Figure 2 and Supplementary Figure 1-3), we explored the expression pattern of tau exons 2 and 3 with and without exon 4a-L across the nervous system (Figure 3d-f). We observed that splicing of *MAPT* exons 2 and 3 leads to expression of 42.6 % 0N, 43.4 % 1N and 13.0 % 2N small tau isoforms in the brain (Figure 3d), similar to what has been previously reported.^27,70^ We observed spinal cord has 32 % 0N, 45 % 1N and 12 % 2N small tau isoforms. The PNS (SN, BP and DRG) contains 37.7 % 0N, 49.2 % 1N and 14.3 % 2N small tau isoforms (Figure 3d).

The big tau isoforms (with exon 4a-L) were present at low abundance in the brain (0.0004 ± 0.0005 0N; 0.002 ± 0.003, 1N and 0.001 ± 0.001, 2N big tau isoforms compared to total-tau) (Figure 3e). Compositionally, we detected 0.03 % 0N, 0.16 % 1N, and 0.08 % 2N big tau isoforms out of total tau detected in the brain (Figure 3e). In contrast, in adult human spinal cord we detected 0.005 ± 0.001 0N, 0.06 ± 0.02 1N and 2N big tau isoforms compared to total tau (Figure 3e). Compositionally, adult spinal cords contain 4 % 0N and 48 % 1N and 48 % 2N big tau isoforms. Further, we also observed that PNS contains 15.0 % 0N (0.086 ± 0.090 compared to total tau), 40 % 1N (0.244 ± 0.190), and 45 % 2N (0.718 ± 0.150) big tau isoforms (Figure 3e). These results highlight the differential splicing of exons 2 and 3 for small and big tau isoforms in the adult nervous system. The exclusion of both exons 2 and 3 is more common for small tau, while favoring a larger N-terminal projection that exclusively contains exons2/3 for “big-tau”. We demonstrated that, while the majority of tau expressed in the CNS, brain (99.7 %) and spinal cord (89.3 %) is small tau (0N+1N+2N small tau isoforms), there is a considerable amount of small tau (∼ 65 % in SN, 50 % in BP and 44 % in DRG) expression in the PNS (Figure 3f). Compositionally, big tau isoforms (0N+1N+2N big tau) are significantly increased to 56 % in DRG (*p* < 0.0001), 50 % in BP (*p* < 0.0001), 35.1 % in the SN (*p* < 0.0001), and 10.7 % in the spinal cord (*p* < 0.0001) compared to 0.3 % in the brain (Figure 3f). The distribution of these big tau 4a-L isoforms further validated the relative expression pattern of tau exon 4a-L from CNS to PNS across the nervous system (Figure 3g).

### Regional distribution of big tau in the human brain

In rodents, big tau expression appears enriched in the cerebellum relative to cortical regions, although prior studies have reported inconsistent cortical expression.^40,55^ To investigate if there is similar regional variance in the human brain, we quantified the distribution of small and big tau peptides from six brain regions. We calculated the total contribution of small (Figure 4a) and big tau (Figure 3b) N-terminal isoforms in these brain regions. The major contribution of small tau isoforms (0N+1N+2N) to the total of tau species remained unaltered across brain cortical regions (temporal, occipital, frontal) and the cerebellum (Figure 4a). Parietal small tau isoform levels (1.48 ± 0.18 normalized to total-tau) were significantly higher than frontal cortex (1.1 ± 0.15) (Figure 4a, *p* < 0.0001). Regarding the minor big tau isoforms (0N+1N+2N), they were significantly increased in the cerebellum (0.018 ± 0.009 normalized to total-tau, *p* < 0.001) compared to temporal (0.005 ± 0.004) and parietal (0.002 ± 0.001) brain regions (Figure 4b). The analysis showed that cerebellum has the highest big tau isoforms (1.2 % of total tau) with decreasing levels in the cortical regions in the following order: frontal cortex (0.4 %) > occipital cortex (0.3 %) > temporal cortex ∼ parietal cortex (0.2%) (Figure 4c). Big tau isoforms remain relatively low in abundance (less than 1 %) in comparison to small tau isoforms across the cortical regions (99 %) and the cerebellum (98 %).

**Figure 4:**
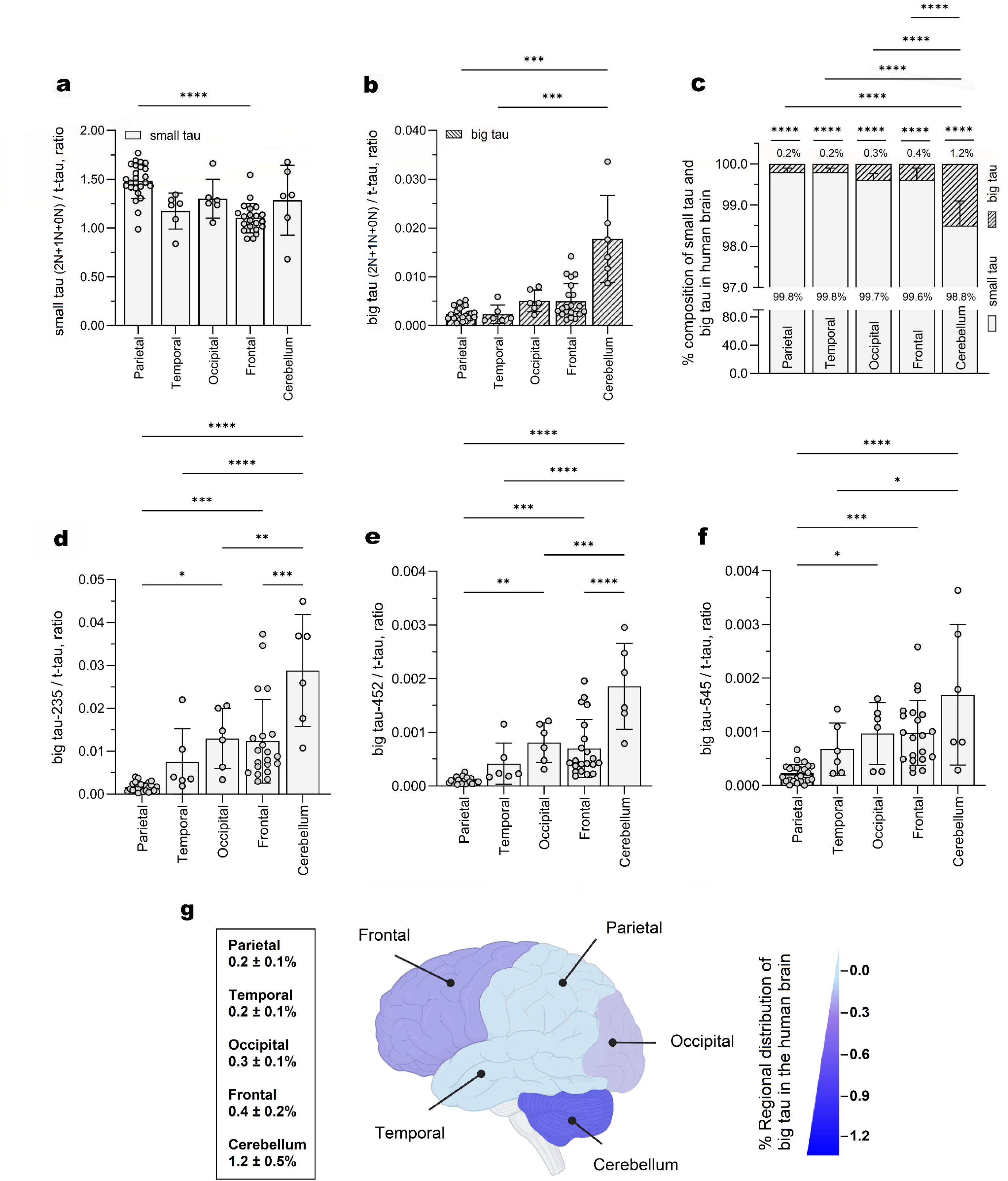
Regional distribution of small and big tau in the adult human brain. **a.** Bar graphs of the relative distribution of the small tau isoforms (0N+1N+2N) normalized to t-tau (residue 633-642) within human brain: parietal cortex (n=25), frontal cortex (n=21), temporal cortex (n=6), occipital cortex (n=6) and cerebellum (n=6). Data are expressed as mean ± SD. Data were analyzed using non-parametric one-way ANOVA followed by Dunn’s multiple comparisons test, ****p < 0.0001, ***p = 0.001. b. Bar graphs of the relative distribution of the big tau isoforms (0N+1N+2N) normalized to t-tau with the human brain regions for all the samples used in (**a**). All values are expressed as mean ± SD. Data were analyzed using non-parametric one-way ANOVA followed by Dunn’s multiple comparisons test, ****p < 0.0001, ***p = 0.001. c. Bar graphs depicting the relative contribution of small tau isoforms and big tau isoforms (0N+1N+2N) across the human brain for all the samples used in (**a**). Data are expressed as mean ± SD. Statistical analysis performed using two-way ANOVA with Tukey’s multiple comparisons test, ****p < 0.0001. **d-f.** Bar graphs of the relative exon 4a-L peptides (residue 235-242 and residue 452-461) and exon 6 peptide (residue 545-552) ratios for all the samples used in (**a**). Data are presented as mean ± SD. Statistical analysis was performed using ordinary one-way ANOVA with Tukey’s multiple comparisons, ****p < 0.0001, ***p ≤ 0.0006, **p ≤ 0.005, *p ≤ 0.0463. **g.** Schematic diagram for the regional distribution in the human brain demonstrates heterogeneity in the expression of big tau protein.

Brain regional distribution pattern for big tau was further validated on other exon 4a-L peptides that were significantly increased in the cerebellum (residue 235-242, 0.03 ± 0.01 and residue 452-461, 0.002 ± 0.001) compared to other cortical regions (*p* < 0.0001) (Figure 4d-f and Supplementary Figure 7). A significant increase was detected for the exon 6 peptide (residue 545-552) in the cerebellum (0.002± 0.001) compared to parietal (0.0002 ± 0.0001, *p* < 0.0001) and temporal cortical regions (0.001 ± 0.0005, *p* = 0.0261) (Figure 4f). These results confirm regional variance of exon 4a-L and exon 6 translation in the human brain with the cerebellum containing higher levels of big tau compared to cerebral cortex, which highlighted a 3-fold increase of big tau isoforms in the cerebellum (1.2 %) compared to 0.2 %-0.4 % detectable in the cortical brain regions (Figure 4g and Supplementary Figure 7).

### Small and Big tau peptides in human CSF

Soluble tau in the CSF primarily consists of small tau species derived from brain with a major truncation around the end of the proline rich region (PRR).^14,57^ Recent investigations of CSF tau species (tau isoform and p-tau) have mainly focused on the small tau isoforms due to their closer association with pathological changes within brain in neurodegenerative disorders, while the contribution of the big tau isoforms in this biofluid and their disease relevance remain unexplored. Tau was immunoprecipitated (Tau-1/HJ8.5/HJ8.7) from human CSF and extracellular small tau and big tau specific peptides (exon 4a-L and exon 6 peptides) were quantified to estimate their corresponding contributions to the total tau concentration in the biofluid (Figure 5a). The CSF concentration of total tau (estimated from peptides comprised of residues 06-23, 45-67, 602-611, 616-630, and 633-642) from young normal control participants ranged from 1.65 to 5.8 ng/mL, and that of the truncated tau containing microtubule binding region of tau (MTBR-tau-243, estimated from a peptide spanning residues 243-254) was 0.1-0.2 ng/mL (Supplementary Figure 8).^14^ The CSF concentration of big tau from young normal controls (estimated from peptides derived from exon 4a-L, comprised of residues 235-242, 294-308, 379-395, 429-442, 452-461) ranged from 0.04-0.1 fmol/mL (Figure 5a [blue highlighted] and Supplementary Figure 8), representing ∼ 1 % of total soluble tau (total-tau peptide [30.0 ± 16.9 fmol/mL]) in the CSF, which is comparable to its proportion estimated in the brain (0.3-0.5 % big tau compared to total tau) (Figure 3). The exon 6 peptide (spanning residues 545-552) was detected at even lower levels (0.2 % compared to total tau) (Figure 5a, red highlighted). In contrast, the small tau specific peptide (small tau 0N isoform) was detected at relatively higher levels (0.25 ± 0.01 normalized to total-tau [Figure 5a, green highlighted]) compared to total tau (T212 peptide according to small tau, spanning residues 633-642 in big tau [and as represented in Figure 5a]). This analysis confirms CSF tau isoform distribution is essentially identical to that of the adult human brain.

**Figure 5:**
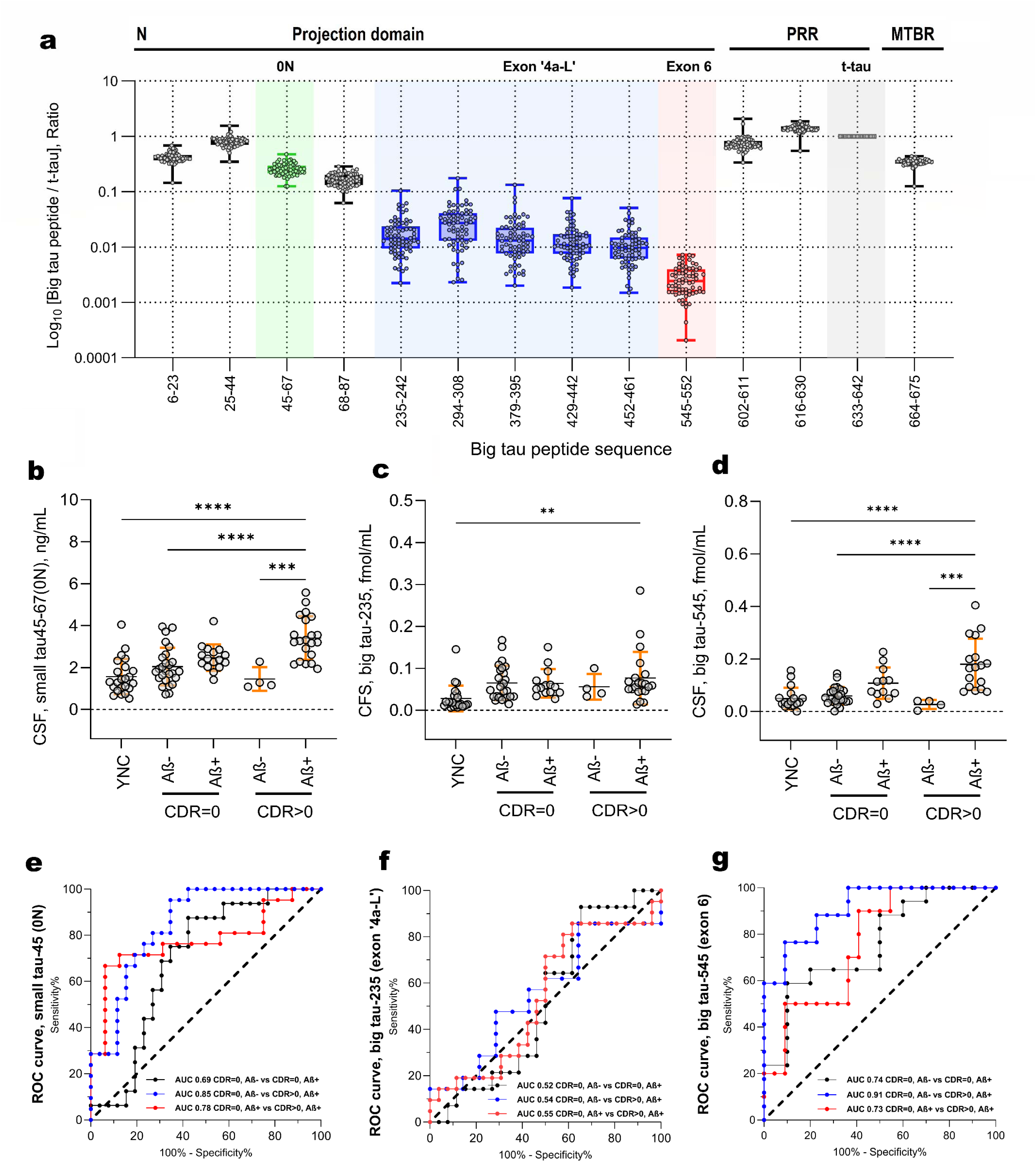
Distribution profile of the small tau and big tau peptides in the human CSF and their respective diagnostic performance as predictive biomarkers in AD continuum. **a.** Box plots of log_10_ normalized CSF big tau profile (n=70; includes young normal controls, Aβ-, Aβ+) quantified using HJ8.5/HJ8.7 and Tau 1 immunoprecipitation focusing on N-terminal tau peptides to mid-domain of tau. CSF small tau 0N isoform peptide (residue 45-67), exon 4a-L, exon 6 peptides and total-tau peptide (t-tau, residue 633-642) is highlighted from left to right. Scatter plots of CSF tau (**b**) small tau 0N isoform (residue 45-67), (**c**) exon 4a-L (residue 235-242) and (**d**) exon 6 (residue 545-552) peptide concentrations from a cross-sectional cohort of young normal controls (n=23), amyloid-negative CDR = 0 (age matched control, n= 26), amyloid-positive CDR = 0 (preclinical AD, n = 14), amyloid-negative CDR > 0 (non-AD with cognitive impairment, n = 4) and amyloid-positive CDR > 0 (symptomatic AD, n = 21). Data are presented as mean ± SD. Statistical analysis conducted by using ordinary one-way ANOVA with multiple comparisons, ****p < 0.0001, ***p = 0.0001, **p = 0.0038. The receiver operating characteristic (ROC) curves and area under the curve (AUC) values for (**e**) small tau 0N (residue 45-67) isoform, (**f**) exon 4a-L (residue 235-242) and (**g**) exon 6 (residue 545-552) peptides demonstrating their respective differential diagnostic accuracies in differentiating symptomatic clinical stage of AD compared to preclinical AD and age-matched controls.

### CSF small and big tau concentrations in AD cross-sectional cohort

To begin to evaluate if small tau and big tau (exon 4a-L/exon 6) peptides in CSF are changed in AD, we examined the CSF from a cross-sectional cohort^57^ of amyloid-negative and amyloid-positive participants representing different clinical stages: young normal controls (n=23), older amyloid-negative CDR = 0 participants (age matched control, n= 26), older amyloid-positive CDR = 0 participants (preclinical AD, n = 14), older amyloid-negative CDR > 0 participants (non-AD dementia, n = 4) and older amyloid-positive CDR > 0 participants (symptomatic AD, n = 21). The CSF small tau levels (0N isoform) showed statistically significant differences between symptomatic AD (CDR > 0, Aβ+) and age-matched controls (CDR = 0, Aβ-) (*p* = 0.0008) as well as preclinical AD (CDR = 0, Aβ+) (*p* = 0.014) and non-AD dementia (Figure 5b). CSF big tau exon 4a-L tau peptide levels (e.g., that spanning residues 235-242) were only significantly increased in symptomatic AD (CDR > 0, Aβ+) individuals compared to young normal controls (YNC) (*p* < 0.0001) (Figure 5c) and not in comparison to age-matched controls or across the AD stages. CSF levels of the exon 6 peptide (residues 545-552) were significantly increased in symptomatic AD individuals compared to young and age matched controls (*p* < 0.0001) and those with non-AD dementia (*p* = 0.03) (Figure 5d), while no significant difference was observed for exon 6 peptide between preclinical AD and symptomatic AD.

Next, we investigated the correlation of CSF small and big tau species with other CSF tau species such as p-tau and truncated tau. We quantified multiple tau species (non-phosphorylated tau peptides comprised of residues 06-23, 25-44, 68-87, 45-67 [0N small tau], 602-611, 633-642), phospho-tau (phospho-tau pT181 and pT217), big tau (peptide comprised of residues 235-242, 2294-308, 429-437 and 452-461) and MTBR-tau243 in the CSF from these participants (Supplementary Figures 8-9) and correlated them with small and big tau levels across the whole cohort (Supplementary Figure 10). The small tau (0N isoform) peptide strongly correlated with other common tau species (e.g., ρ = 0.93, *p* < 0.0001 with total-tau (T212-221); and ρ = 0.97, *p* < 0.0001 with total-tau (T181-190)) (Supplementary Figure 10), correlated weakly with phospho-tau pT181 (ρ = 0.44, *p* < 0.0001) and phospho-tau pT217 (ρ = 0.51, *p* < 0.0001 with pT217/T217) (Supplementary Figure 10), a biomarker that is highly correlated with amyloidosis in preclinical AD, ^71,72^ and correlated strongly with CSF MTBR-tau243 (ρ = 0.87, *p* < 0.0001) (Supplementary Figure 10), a truncated tau species in CSF that exists at baseline in preclinical AD and has been demonstrated to increase and correlate with NFT pathology in AD.^14,73^ This highlights that small tau 0N isoform starts to increase when individuals that are positive for amyloid become symptomatic, in contrast to phospho-tau species that increase when individuals become positive for amyloid, and that have therefore become established biomarkers for preclinical (asymptomatic) AD. We further investigated the association of clinical cognitive measures (MMSE) and CSF biomarkers (Supplementary Figure 11). As previously demonstrated, CSF MTBR-tau243, a biomarker of AD tau tangle pathology, was negatively correlated with cognitive performance in our cohort (ρ = -0.45, p = 0.0012; Supplementary Figure 11A), with higher levels associated with poorer cognitive outcomes.^15^ Likewise, N-terminal tau (represented by a peptide spanning residues 06-23), small tau (0N isoform) and total-tau (represented by a peptide spanning residues 181-190) demonstrated significant negative associations with poorer cognitive outcomes (Supplementary Figure 11B-D, F). Importantly, we found that percent occupancy of phosphorylation at pT217 (pT217/T217 %) also demonstrated a strong negative correlation with cognitive outcomes (ρ = -0.52, *p <* 0.0001, Supplementary Figure 11G), whereas the percent pT181 (pT181/T181 %) did not demonstrate a significant association with cognitive outcomes (ρ = -0.11, *p* = 0.44, Supplementary Figure 11G). In contrast, the CSF big tau exon 4a-L peptide (residue 235-242) weakly correlated with total tau (T212; ρ = 0.34, *p* = 0.02), pT217/T217 % (ρ = 0.32) and MTBR-tau243 (ρ = 0.38) (Supplementary Figure 10). Additionally, we observed weak correlations of CSF big tau exon 4a-L peptides (big tau-294, big tau-429, big tau-452) with CSF total tau (T212, ρ = 0.27-0.35), pT217/T217 % (ρ = 0.3) and MTBR-tau243 (ρ = 0.4). These results suggest that CSF levels of soluble exon 4a-L tau species are not impacted by amyloid status (inferred from pT217/T217 %) or tau pathology (CSF MTBR-tau243 and total tau) in AD. In contrast, the exon 6 peptide weakly correlated with CSF tau species (ρ = 0.50, *p* < 0.0001 with total-tau and r = 0.55, *p* < 0.0001 with MTBR-tau243) (Supplementary Figure 10). Additionally, CSF big tau exon 4a-L peptide levels (residues 235-242 and 429-442) demonstrated no correlation with cognitive measures (big tau 235-242; ρ = 0.052, *p* = 0.72 and big tau 429-442; ρ =-0.013, *p* = 0.93; Supplementary Figure 11 H-I), whereas exon 6 peptide (residue 545-552) demonstrated significant negative correlations with cognitive performance (ρ = -0.46, *p* < 0.001, Supplementary Figure 11J). These findings suggest that CSF exon 6 levels change in symptomatic AD and potentially reflect altered tau tangle pathology.

Additionally, we investigated the diagnostic accuracies of the small tau and big tau (exons 4a-L/6) peptides in the CSF when distinguishing individuals at different stages of AD (Figure 5e-g). CSF small tau levels discriminated between CDR > 0, Aβ+ and CDR = 0, Aβ+ with an area under the curve (AUC) [95 % CI] = 0.78 [0.62-0.93] (Figure 5e, red). When discriminating between CDR > 0, Aβ+ and CDR = 0, Aβ-individuals, CSF small tau had an AUC [95 % CI] = 0.85 [0.74-0.96] (Figure 5e, blue). In comparison, CSF big tau peptide (residues 235-242, from exon ‘4a-L’) showed essentially no utility for discriminating CDR > 0, Aβ+ from CDR = 0, Aβ- with an AUC [95 % CI] = 0.54 (Figure 5f, blue) or CDR > 0, Aβ+ and CDR = 0, Aβ+ (Figure 5f, red). In contrast, CSF exon 6 peptide (residues 545-552) discriminated between CDR > 0, Aβ+ and CDR = 0, Aβ+ groups with an AUC [95 % CI] = 0.73 [0.57-0.95] (Figure 5g, red) and between CDR = 0, Aβ- and CDR > 0, Aβ+ groups with an AUC [95 % CI] = 0.91 [0.83-0.99] (Figure 5g, blue).

Finally, we investigated the relationships of small and big tau CSF levels with age for the entire cross-sectional cohort with and without amyloidosis (Figure 6). We observed that the CSF small tau levels were significantly increased for amyloid positive individuals at age > 70 compared to amyloid negative individuals at age > 70 (*p* = 0.02) and amyloid negative individuals in the age group 40-60 (*p* = 0.001) (Figure 6a and Supplementary Figure 10). CSF exon 4a-L big tau peptide increased with age and was significantly increased in amyloid positive individuals in the age group > 70 compared to normal controls in the age group 19-40 (*p* = 0.04) (Figure 6b). CSF exon 6 peptide was significantly increased in amyloid positive individuals in the age group > 70 compared to amyloid negative individuals in the age group > 70 (*p* < 0.001), normal controls in the age group 19-40 (*p* = 0.01) and amyloid negative individuals in the age group 60-70 (*p* = 0.04) (Figure 6c). In amyloid negative individuals, we observed big tau peptide (residue 235-242) correlated better with age (r = 0.56, *p* < 0.0001) compared to MTBR-tau243 (r = 0.47, *p* < 0.0001), small tau (r =0.3) and total-tau (r =0.36, *p* < 0.0001) (Figure 6d).

**Figure 6:**
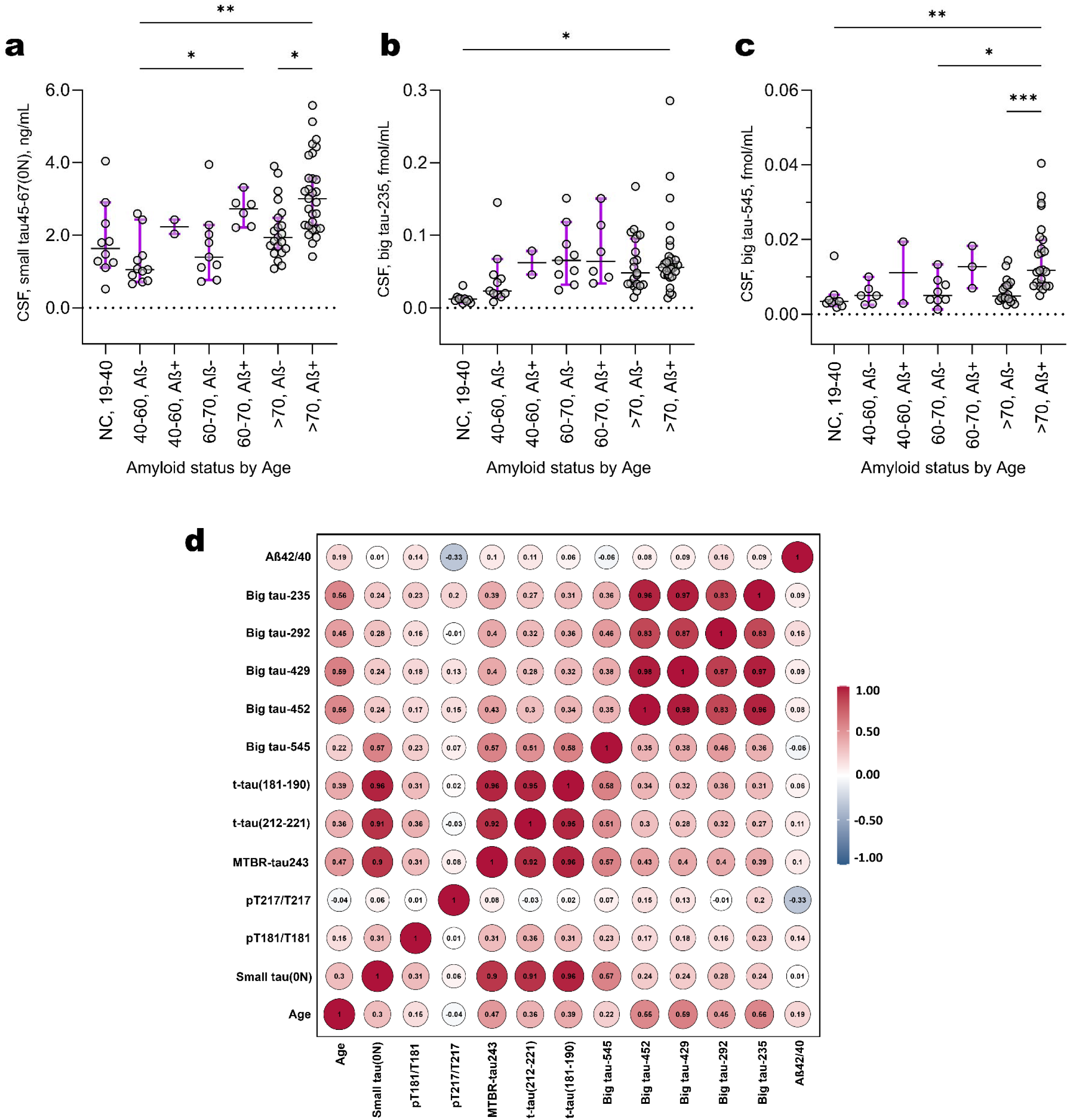
CSF exon 4a-L peptides increase with age independently of cerebral amyloidosis. Scatter plots of CSF tau (a) small tau 0N isoform, (b) exon 4a-L (residue 235-242) and (c) exon 6 peptide concentrations from a cross-sectional cohort in different age groups with and without amyloidosis (Aβ-, n=41; Aβ+, n=37). Data are presented as mean ± SD. Statistical analysis was performed using two-way ANOVA with multiple comparisons, ***p = 0.0006, **p ≤ 0.0088. d. Correlation matrix presenting Spearman’s correlation for all the tau biomarkers with each other in only Aβ-participants.

In summary, CSF small tau isoform (0N isoform) exhibited changes later in the symptomatic stage of AD, associating better with CSF total-tau as well as CSF MTBR-tau243. CSF big tau exon 6 peptide also demonstrated changes associated with AD that will need further exploration in a larger cohort, whereas CSF big tau peptides (exon 4a-L) did not change with amyloidosis, tau pathology (CSF MTBR-tau243) or cognitive symptom onset (Supplementary Figures 11-13).

## Discussion

Despite the knowledge of the tau mRNA transcript for the last three decades, the protein distribution of the longest isoform of tau – “big tau” has not been actively investigated in the human nervous system.^40,47,50,65^ Previous qualitative reports have revealed big tau is preferentially expressed in the soma, dendrites and axons of the CNS neurons that extend processes into the periphery, in addition to PNS neurons and optic nerves.^40^ Exon 4a-long has been previously reported in human cancer cells and skeletal muscle tissues (non-neuronal tissues) as the primary constituent of HMW tau along with the common small tau isoforms.^45,46^ This larger splice variant, exon 4a-L (1065 base pairs (bp)) was found to be expressed in higher amounts (by RT-PCR) than the 753 bp exon-4a-S.^46^ The exon 4a-L contains two 3’ splice sites and when the alternative 3’ splice junction (CAG/A) within this exon is utilized it results in the insertion of an additional 312 bp between exons 4 and exon 4a-S.^45^ Using tandem mass spectrometry sequencing techniques, here we report that this longer exon 4a (exon 4a-L) is predominantly expressed across the human nervous system and encodes for 355 amino acids. This longer exon 4a is inserted between tau exons 4 and 5 and has an additional 104 amino acids compared to the exon 4a-S (251 amino acids).^50,55^ Tandem mass spectrometry using protein database searching led to the discovery of the exon 4 to exon 4a-L junction/bridging peptides in human nervous system tissues. The alternative 3’ splice site resulting in exon 4a-L has been proposed as favored splice product due to the presence of consensus (i) 3’ splice junction, (ii) branch point sequence (BPS) upstream in the genomic sequence and (iii) polypyrimidine rich sequence (28 bp) following the BPS. Long read sequencing data from control brain samples (caudate and middle frontal gyrus) further confirmed that the prominent form of big tau in the CNS is the exon 4a-L isoform. Low transcript levels of exon 4a-L isoforms in the selected brain regions were consistent with the quantitative proteomic results of low relative abundances of big tau in the brain (< 0.5 % of total tau). Future long-read sequencing across additional brain regions (cerebrum, cerebellum, and brainstem) will help define the anatomical heterogeneity of big tau isoform expression (4a-L vs. 4a-S). Targeted mass spectrometry using synthetic standard peptides for big tau exon 4a-L and exon 4a-S isoforms demonstrated that exon 4a-L is the predominant form (> 95 % of big tau) in human nervous tissues, whereas exon 4a-S isoforms constitutes a minor fraction (< 5% of big tau). Notably, exon 4a-S was relatively more abundant in the PNS (∼3.5% of big tau) than in the CNS (< 1% of big tau), although exon 4a-L remained the dominant isoform across regions. Altogether, our results expand the repertoire of human big tau isoforms beyond what has been previously reported in the literature.

To complement the mass spectrometry and transcriptomic analyses, we employed immunoblotting to resolve tau species by molecular weight, enabling detection of isoform heterogeneity and high molecular weight (HMW) species not readily distinguished by bulk RNA sequencing or peptide-based proteomics. We observed a range of HMW bands ∼ 115-90 kDa in PNS and spinal cord tissue samples on immunoblots using Tau-1 and tau-5 antibodies, while no discernable HMW bands were detected in the brain regions investigated. Tau PTMs (truncation, phosphorylation, acetylation etc.) could result in multiple HMW bands of big tau. Future investigations using big tau specific antibodies on different tissue lysates will provide further insights on the origin of multiple HMW tau bands beyond the alternative spliced big tau isoforms (exon 4a-L/4a-S). SDS-PAGE followed by in-gel digestion and LC-MS of these HMW bands from CNS and PNS tissue lysates confirmed the presence of exon 4-4a-L junction peptides. The insertion and expression of exon 4a-L along with exon 6 results in an isoform of tau (big tau) that is almost twice as long (862 amino acids) as the conventional small tau isoform (441 amino acids). The exon 4a between human and rodent nervous system are different. The protein sequences of big tau (exon 4a) in mouse and rat retain only 50 % and 51 % homology, respectively, with human big tau.^50^ Another between-species distinction of big tau sequence could be explained by the presence of an alternative 3’ splice site within human tau genome. In rodents, there is a stop codon within the additional 312 bp sequence and their genomic sequence lacks consensus BPS analogous to a human branchpoint. Thus, while our results indicate that human big tau is primarily derived from exon 4a-L insertion, we expect rodents will primarily express the smaller HMW tau isoform (exon 4a-S). The expression pattern of exon 4a-L and exon 4a-S in rodents needs further investigation.

Ever since its discovery, several descriptive reports have tried to delineate the distribution of big tau in the PNS and optic nerves along with some selected CNS regions.^40,47,65^ In this study, we found relative differential distribution pattern of exon 4a-L expression increasing from CNS to PNS across the human nervous system in the following order: brain << spinal cord < sciatic nerve < brachial plexus ∼ dorsal root ganglion. More importantly, while small tau is the major tau isoform in the CNS (∼ 99 % in brain and ∼ 90 % in spinal cord), our results indicate there are almost equimolar amounts of small tau and big tau in the PNS (∼ 44-55 %). Our results challenge the idea that small tau is CNS specific with big tau being exclusive to the PNS. Instead, we propose small tau being CNS enriched, while the PNS is enriched with big tau isoforms. Our results also highlight that while the percentage of big tau contribution increases from CNS to PNS, the total tau concentration from CNS to PNS decreases by ∼3-orders of magnitude. We found the brain (∼ 10 ng/µg protein) has almost 1000-fold higher total tau levels than the peripheral nerves (∼ 0.03 ng/µg protein). Similar distribution of total tau has been previously reported, where peripheral tissues demonstrated ∼ 1.8 % (submandibular gland) to 0.16 % (liver) total tau compared to the brain (∼ 7800 ng/mg brain).^41^ Another important finding of our study was the relatively low exon 6 peptide abundance in spinal cord and PNS, indicating exon 6 is weakly expressed in these tissues. This finding could potentially explain why big tau isoforms are different in neural cell lines compared to DRG, spinal cord and sciatic nerve, which selectively express the exon 4a-L and not exon 6.^39^ Inclusion of exon 6 in both the 6 and 9 kb tau mRNA and its presence has been reported in both fetal and adult human tau mRNA.^39,67,74^ Exon 6 has also been found in mature and immature spinal cord but not in the PNS, decoupled from the peripheral exon 4a-L.^74^ This could lead to a further increase in the repertoire of tau isoforms beyond what we have reported in this study. Our results indicate exon 6 and exon 4a-L could be mutually exclusive, and splicing could be regulated in tissue-specific manner, something that will need further validation using specific antibodies to tau exon 4a-L and exon 6. Understanding the functional consequences of these differential expression profiles requires further investigation.

Developmental studies of tau expression in human brains have shown three major isoforms of tau at early postnatal stages, versus six isoforms of tau in adulthood.^2,27,28,67,75^ In contrast, big tau isoform stoichiometry and distribution have not been studied until now. Intuitively, cassette splicing of exons 2 and 3 (E2 and E3) should give rise to big tau isoforms with 0, 1 or 2 N-terminal repeats, as has been observed for small tau. Our characterization of the exon 4a-L sequence was crucial in identifying the specific exon 4 to exon 4a-L junction in big tau, which led to the discovery of the three distinct N-terminal isoforms for big tau (0N-, 1N-, 2N-big tau). Given that 3R and 4R isoforms were also quite abundant in the PNS, we speculate that E10 splicing for big tau would be a common feature, resulting in six distinct isoforms of big tau, similar to the six isoforms of small tau. Consistent with previous reports,^27,68^ we observed adult human brains have 43 % 0N, 46 % 1N and 13 % 2N small tau isoforms, with subtle increase in 1N small tau isoform across spinal cord to PNS. Unlike small tau isoforms, we discovered adult human nervous tissue contains equimolar levels of 1N and 2N (40-50 %) big tau isoforms, with 0N big tau isoform being present at less than 5 %. These observations, based on multiple CNS (brain and spinal cord) and PNS (sciatic nerve) tissue lysates from cognitively normal control individuals, and individuals with AD and ALS, highlighted that the extended N-terminal projection domain in big tau is a common feature across the adult human nervous system. Alternatively spliced isoforms of the same protein can have distinct biological functions, with some alternative isoforms being functionally divergent.^76^ While the specific physiological roles of big tau remain unclear, the “long” exon 4a insertion is hypothesized to significantly increase the spacing between microtubules, compared to small tau. Similar dramatic structural changes have also been associated with other microtubule associated proteins (MAP); the low molecular weight isoform (∼ 70 kDa) of MAP2C (juvenile form of MAP2) switches to higher molecular weight MAP2A/B (∼ 250 kDa) during later stages of neuronal development.^77,78^ The N-terminal projection domain of tau, which functions as a variable spacer between neighboring microtubules and regulated by alternative splicing of E2/E3, may play a developmental role in modulating inter-microtubule spacing.^79,80^ Indeed, cross-sections of axonal microtubules reveal a larger spacing of ∼35 nm when induced by big tau compared with ∼ 20 nm when induced by small tau isoforms in neuronal cultures.^80^ This larger spacing has been suggested to provide more efficient organelle transport in the axon by lowering resistance of the axoplasm compared to the small tau, that is more conducive to axonal growth.^47^ The presence of big tau in the long and high caliber neurons in the periphery may be driven by the need for robust and energy efficient axonal transport, while small tau isoforms provide more plasticity to the CNS neurons. Developmental studies on big tau expression in rodents indicate active translation in postnatal stages,^40,50^ and further exploration would provide clues why we observe a distinct dissociation in relative isoform distribution between small and big tau.

Selective vulnerability is a common phenomenon in many neurodegenerative disorders that feature distinct patterns of neuronal loss and accumulation of protein aggregates within certain brain regions, while other regions are resistant to the pathology.^10,81–83^ Stereotactic spread of tau pathology (neurofibrillary tangles) in AD and other tauopathies^11,84,85^ is thought to occur throughout the hippocampus and the cortex, sparing very few brain regions such as cerebellar cortex even in extreme cases.^10,86^ Tau expression is ubiquitous within the brain, while the selective vulnerability of certain brain regions in tauopathy remains an intriguing outstanding question. While prior studies have examined expression and splicing regulation of tau across different brain regions,^28,68^ currently the exact reasons for this regional vulnerability are not yet known. Interestingly, exon 3-encoded inserts have been shown to inhibit tau aggregation, while exon 2 and 10-encoded inserts increase the aggregation propensity of tau.^87^ Recent research has highlighted that big tau has significantly diminished aggregation propensity compared to the small tau isoforms.^88^ They found that exon 4a (exon 4a-S) has enhanced microtubule-binding capacity and exhibited less hyperphosphorylation – key mechanisms that ultimately promote tau aggregation.^3,89^ Future investigations are needed to determine whether exon 4a-L big tau isoforms similarly exhibit enhanced microtubule binding along with reduced aggregation propensity. Indeed, in our bottom-up proteomic investigation across the nervous system, we did not observe evidence for phosphorylated exon 4a peptides. Moreover, multiple mutations and polymorphisms have been identified on the exon 4a in humans without any known pathogenicity.^90–93^ Our finding of 3-fold increase in exon 4a-L expression in the cerebellum compared to other cortical brain regions aligns with previous reports in human brains and rodents,^40,47,88^ emphasizing the regional heterogeneity of big tau expression in the brain. Most interestingly, the lowest levels of exon 4a-L in the brain coincide with the vulnerable regions for AD. This raises the intriguing prospect of therapeutically altering the small and big tau ratios in these brain regions to recruit the neuroprotective role of big tau in resisting tangle pathology.^55,94^

CSF is thought to closely reflect brain pathological processes, carrying molecules through passive diffusion from the brain parenchyma. In this study, we finally investigated CSF to understand the tau isoforms distribution in this biofluid and their relation to AD. This is the first study to quantitively measure and compare the small tau and big tau peptides in human CSF. Normalizing the small tau (0N isoform) peptide in the CSF to the total tau peptide indicated this isoform is quite abundant, whereas big tau (exon 4a-L/exon 6) accounts for less than 1 % of total tau in the CSF. This is consistent with the relative abundance of exon 4a-L and exon 6 in the brain. CSF total-tau (t-tau) measured by monoclonal antibodies binding to the mid-domain of tau is known to increase during acute neurodegeneration.^95,96^ Assays that measure endogenous brain-derived fragments of tau species from the N-terminus and MTBR have been developed to improve the diagnosis of AD and other tauopathies.^14,19,73^ Recent studies highlight the importance of brain-derived tau species in the CSF/plasma as neurodegeneration biomarkers useful in identifying and monitoring neurodegenerative progression in AD.^16,19,20^ Plasma N-terminal containing tau fragments (NTA-tau) and brain-derived tau (BD-tau) assays were first validated from cross-sectional AD cohorts with paired CSF and plasma samples from cognitively normal and symptomatic individuals. Our mass spectrometry data provides plausible tau species that are also targeted by these assays for detecting tau pathology in AD. In this study, we found significant increase of CSF small tau (0N isoform) in amyloid positive symptomatic individuals (CDR > 0). High correlation of the CSF small tau (0N isoform) with total-tau and MTBR-tau243, and not so much with p-tau measures (pT181/T181 and pT217/T217), suggests a closer link to the clinical phase of AD.^14,17,72^ In contrast, we found CSF big tau exon 4a-L increased with age but no significant changes of the CSF big tau exon 4a-L peptides in either preclinical AD or in individuals with mild cognitive impairment due to AD. We also compared our results with CSF MTBR-tau243, a high performing fluid biomarker specific for AD tangle pathology. CSF big tau exon 4a-L peptides poorly correlated with total-tau biomarkers and MTBR-tau243 (ρ=0.38). Our results thus identify the small tau (0N isoform) as an important biomarker target in the blood similar to brain-derived tau^16,20^ that might function as a neurodegeneration biomarker in AD. Translation of this IP-MS assay, that can simultaneously quantify small and big tau isoforms, into blood derived plasma would supplement current plasma BD-tau and NTA-tau assays in disease monitoring and staging. Interestingly, exon 6 peptide had better clinical performance compared to exon 4a-L peptides, highlighting the complex nature of the CSF tau fragments. Exon 6 peptide demonstrated better correlation with worsening cognitive outcomes (Supplementary Figure 10) as well as CSF MTBR-tau243 (ρ = 0.55). Tissue-specific exon 6 splicing independent of other exons (i.e. exon 4a) has been documented.^97^ The exact mechanism of tau exon 6 splicing, independent of exon 4a-L, and how that correlates with tau pathology in AD requires further investigation.

The enrichment of big tau isoforms in the PNS compared to the CNS suggests that it may be leveraged as a potential biomarker for peripheral nervous system injury, such as peripheral neuropathies. Currently there are no assays that can simultaneously distinguish between the CNS and PNS enriched forms of tau. Recent studies have indicated that plasma p-tau (p-T181) is elevated in sporadic ALS patients with predominant lower motor neuron involvement.^22^ A recent multi-center study assessing serum and plasma p-tau levels in ALS participants has raised concern about the specificity of the p-tau assays. Interestingly, they found elevation of plasma p-tau 181 and p-tau 217 levels in ALS patients, overlapping with levels seen in clinically confirmed AD cases.^23^ Furthermore, immunohistochemical analysis revealed significantly elevated sarcoplasmic reactivity for p-tau 181 and 217 in ALS muscle biopsies compared to disease controls. However, these existing assays cannot differentiate between the phosphorylated tau that could be derived from small tau and big tau isoforms. Such distinction of tau isoform would be crucial in discerning their pathophysiological roles in ALS and other pathologies that impact the lower motor neurons and skeletal muscles. Tau hyperphosphorylation has also been documented in retinal tissues from AD and primary tauopathy participants.^98^ Another report has highlighted tangle like aggregates of hyperphosphorylated tau in the heart tissue of patients with heart failure (HF) and AD, where the predominant isoform of tau is big tau.^66^ While these studies investigated the role of HMW tau, they did not demonstrate whether this big tau protein originated form the insertion from either exon 4a-L or exon 4a-S between exons 4 and 5. Our assay thus provides a unique opportunity to probe disease specificity and investigate what role these divergent tau isoforms play in the pathophysiology of neurodegeneration across the nervous system where tau is implicated.

While our study provides valuable new insights into the fundamental biology of big tau and isoform distribution across the human nervous system, several limitations should be acknowledged. First, our mass spectrometry results are derived from bulk tissue soluble homogenates. While appropriate for peptide sequencing (especially for characterizing the exon junction peptides) as well as quantitative profiling, this method has limitations regarding the spatial and cellular distribution of the big tau across the brain and spinal cord (grey and white matter). Future spatial localization using immunohistochemistry or other spatial omics technologies would be useful to study the functional role of big tau isoforms. Development of human big tau specific antibodies would be useful to confirm the presence of the big tau protein isoforms across different brain regions (cerebrum, brain stem and cerebellum) as well as its specific cellular expression pattern. Using PCR to conclusively confirm the cellular origin of big tau/4a-L and 4a-S in different brain regions will be highly complementary to anatomical profiling. Secondly, the lack of functional studies exploring the biological roles of exon 4a-L/4a-S limits the comprehensive understanding of big tau and its significance in neurodegenerative processes. Thirdly, we used complimentary proteomic tools to investigate the insertion of exon 4a-L/4a-S in the HMW human tau isoform, however our proteomic digestion limits the investigation of simultaneous exon expression (whether exon 4a-L/4a-S and exon 6 are mutually exclusive). Complimentary tools such as long-read sequencing and size-exclusion chromatography along with intact/top-down mass spectrometry could be used to address this in future research. Lastly, while this study evaluates the diagnostic potential of small and big tau species in the CSF in a cross sectional cohort addressing Aβ and tau pathology, further research correlating these distinct tau proteoforms in diverse neuropathological conditions, clinical parameters and disease progression (tau pathology) is necessary to enhance their clinical relevance and utility in disease management. A more extensive analysis of CSF and plasma big tau isoforms in diseases that involve peripheral tau changes, such as peripheral nerve injury and lower motor neuron injury, should be investigated to assess its diagnostic potential.

In summary, here we investigated the tau isoforms with and without exon 4a in human nervous system. Canonical tau isoforms lacking exon 4a account for 99 % of the total tau in the human brain. Insertion of exon 4a between exons 4 and 5 generates a larger tau isoform (big tau), that is twice as large compared to common tau (small tau). Exon 4a undergoes alternative 3′ splicing to generate big tau 4a-L isoforms along with big tau 4a-S isoforms. We demonstrate through the detection of exon-bridging peptides that exon 4a-L is the predominant splice product in humans. Transcriptomic investigation (Isoseq) from two different brain regions identified big tau isoforms with exon 4a-L. Big tau isoforms containing exon 4a-L display a striking regional distribution, comprising ∼1% of total tau in brain but up to ∼60% in the PNS. Alternative splicing of exons 2 and 3 further expands big tau diversity, generating multiple isoforms with distribution patterns distinct from the well-characterized small tau isoforms (0N, 1N, and 2N). Notably, increased expression of big tau isoforms coincides with the brain regions that are resistant to tau aggregation, which raises an intriguing question of whether these tau isoforms are protective against tauopathies and requires further investigation. More importantly, our results demonstrate that, whereas brain derived small tau isoforms in the CSF are increased in AD, CSF big tau peptides remain unaltered. These results provide potential targets for future blood-based biomarker assay development capable of distinguishing between brain derived tau and peripheral tau, which may be crucial for differentiating CNS and PNS diseases that may coincide or overlap in older populations. Together, our findings expand the known tau isoform repertoire and establish a framework for studying CNS- and PNS-enriched tau species in health and disease.

## Acknowledgement

We thank the participants and families for their generous donation of biosamples. This research was supported by Tracy Family Stable Isotope Labeling Quantitation Center established by the Tracy Family, Richard Frimel & Gary Werths, GHR Foundation, Pat and Jane Tracy, Anonymous, Anne & Ray Capestrain, Community Foundation Serving West Central Illinois and Northeast Missouri, JTL Family Fund, Payne Family, Mary & Jay Sullivan, Tracy Family Foundation, Catherine & Tom Tracy, Community Foundation for the Land of Lincoln, Jim & Jil Tracy, Joe & Jill Tracy, Sonja & Robert M. Willman, Boniface Foundation, Jean & Michael Buckley, Ann Liberman, Clemence S. Lieber Foundation, Mary Schoolman & Dr. James Hinrichs, and Susan & Scott Stamerjohn brought together by The Foundation for Barnes-Jewish Hospital. This work was supported by funds provided by the McDonnell Center for Cellular and Molecular Neurobiology at Washington University (S.M.), Cure Alzheimer’s Fund (RJB), Coins for Alzheimer’s Research Trust grant (C.S.), Target ALS for Washington University ALS Postmortem Core (C.V.L., R.J.P.), NIH/NINDS R01NS095773 (R.J.B., C.S.), Rainwater Foundation (R.J.B., C.S., C.M.K, N.G., R.W.P), AFTD (R.J.B., C.S., N.G., R.W.P), National Institute of Health (NIH) NS123985 (C.M.K.), NIH NS110890 (C.M.K.), NIH/NIA P01 AG03991 (R.J.P., E.E.F), NIH P41 GM103422, NIH/NIA P30 AG066444 (R.J.P., E.E.F), and NIH/NIA P01 AG026276. The iso-seq analysis was funded through NIH/NINDS 1U54NS123746-01 (A. M. G.) and Clinical and Translational Science Awards (CTSA) grant UL1TR004419 from the National Center for Advancing Translational Sciences. We acknowledge the help of Dr. Brian Gordon, Reid Coyle, Gina Collins, Janice Chang, Dr. James Bollinger Vitaliy Ovod, Dr. Arun Renganathan, Marsh Jacob, Emma Starr, Brunda Tumala for their scientific and editing help across this study. We thank Dr. David Holtzman and Ms. Hong Jiang for HJ antibodies. We thank the participants and personnel of the Charles F. and Joanne Knight Alzheimer Disease Research Center, as well as the staff of Washington University’s Translational Human Neurodegenerative Disease Research (THuNDR) Laboratory for providing postmortem human brain tissue samples for this study. We thank the Human Brain Collection Core (HBCC) at NIMH, the Netherlands Brain Bank (NBB), University of Washington (UWA) and Banner Sun Health Research Institute (BSHRI) for providing the access to brain tissue samples for long read sequencing data, Lea T Grinberg (Human Biology Validation Core) for her help with sample distribution and computational support from Scientific Computing and Data at the Icahn School of Medicine at Mount Sinai for iso-seq data analysis.

## Author Contributions

S.M., C.M.K., N.R.B., C.V.L., C.P. and R.J.B. contributed to the conception and design of the study; R.K.K., N.R.B., K.H., C.V.L., K.F.R., S.K., R.J.P., E.E.F., C.P., J.O., J.M., T.M., C.S., N.G., A. M. G., C.M.K., R.J.B. and S.M. contributed to acquisition and analysis of data; R.K.K., N.R.B., K.H., C.V.L., C.P., J.O., R. J. P., T.M., N.G., C.S., C.M.K., R.J.B. and S.M. contributed to drafting the text or preparing the figures.

## Potential Conflicts of Interest

The authors have nothing to report.

## Data Availability

The mass spectrometry proteomics data have been deposited to the ProteomeXchange Consortium via the PRIDE partner repository^99^ with the dataset identifier PXD059267. Raw iso-seq data files are deposited in GEO: GSE324361. All the data, tau concentrations and CSF biomarker data presented in this study are available from the corresponding author upon reasonable request, and such arrangements are subject to standard data-sharing agreements and approval by the institutional review board.

## Notes

https://www.ebi.ac.uk/pride/PXD059267

### Competing Interest Statement

K.H. is an Eisai-sponsored voluntary research associate professor at Washington University and has received a salary from Eisai. The remaining authors declare no competing interests.

### Summary of Updates

We now have included discovery and validation of big tau with exon 4a-Long within the brain using long-read sequencing data providing transcriptomic evidence. We also have made signifcant changes to validation of big tau isoform validation for both exon 4a-L and exon 4a-S across CNS and PNS. The Figure 2 has been updated to include endogenous compared to synthetic peptide standards along with updated immunoblots. Limitation section has been updated to clarify. Supplemental files updated.

