## Supplementary Figures for "Distribution of big tau isoforms in the human central and peripheral nervous system"

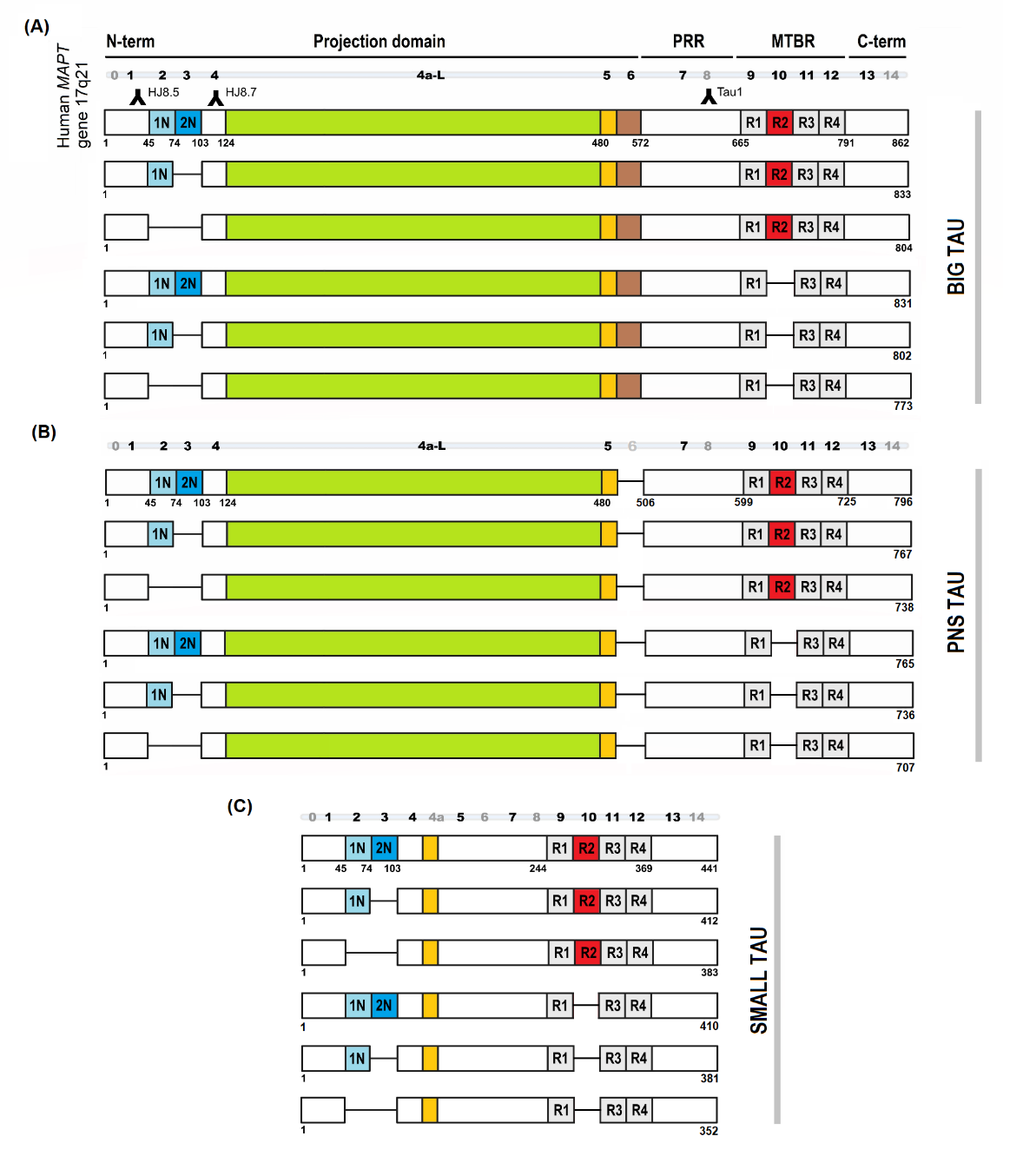
**Supplementary Figure 1a.**

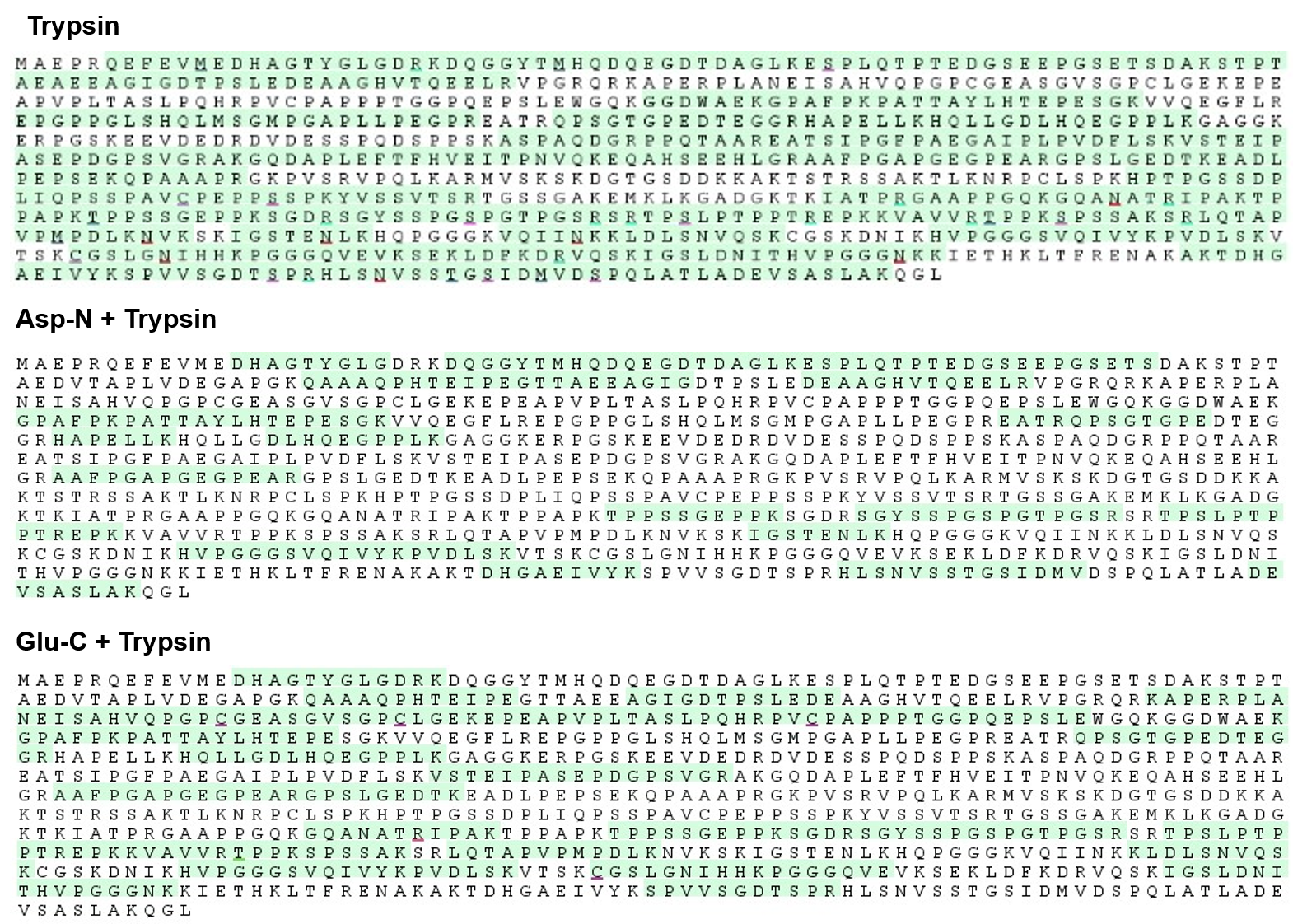
**Supplementary Figure 1b.**

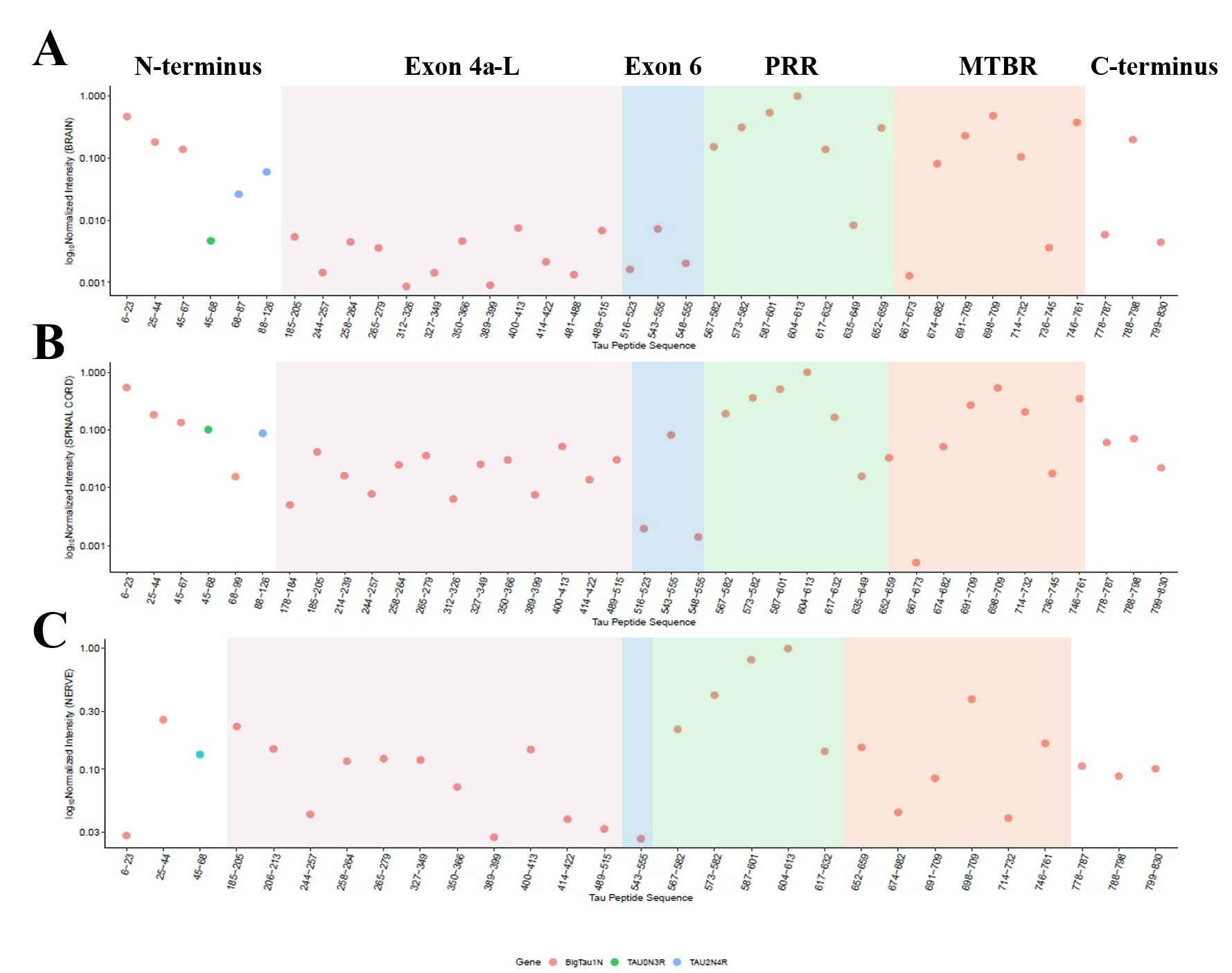
**Supplementary Figure 1c.**

**Supplementary Figure 1a.** Schematic representation of *MAPT* isoforms of (A) Big tau, characterized by the inclusion of exon 4a-L and exon 6, (B) PNS-tau, characterized by the inclusion of exon 4a-L and exclusion of exon 6 and (C) Canonical six CNS tau isoforms (referred to as “small tau”) generated by alternative splicing of exons 2, 3 and 10, comprising both 3R and 4R forms.

**Supplementary Figure 1b.** The protein sequence coverage of tau using bottom-up tandem mass spectrometry (MS/MS) from the human nervous tissue by using combination of enzymatic digestions. The green highlighted regions are the respective peptide identified in the database search in the MSFragger software.

**Supplementary Figure 1c.** The log transformed relative intensity of all the tau peptides along with the exon 4a-L and 6 peptides across the human nervous system derived from bottom-up LC-MS/MS. Compared to (A) brain, the relative intensity of exon 4a-L are two-magnitude higher in the (B) spinal cord and (C) sciatic nerve samples, respectively. The tau peptide sequences are numbered based on the big tau with one N-terminal insert (1N big tau) and ordered from N-terminus to C-terminus (left to right). PRR, proline rich region; MTBR, microtubule binding region.

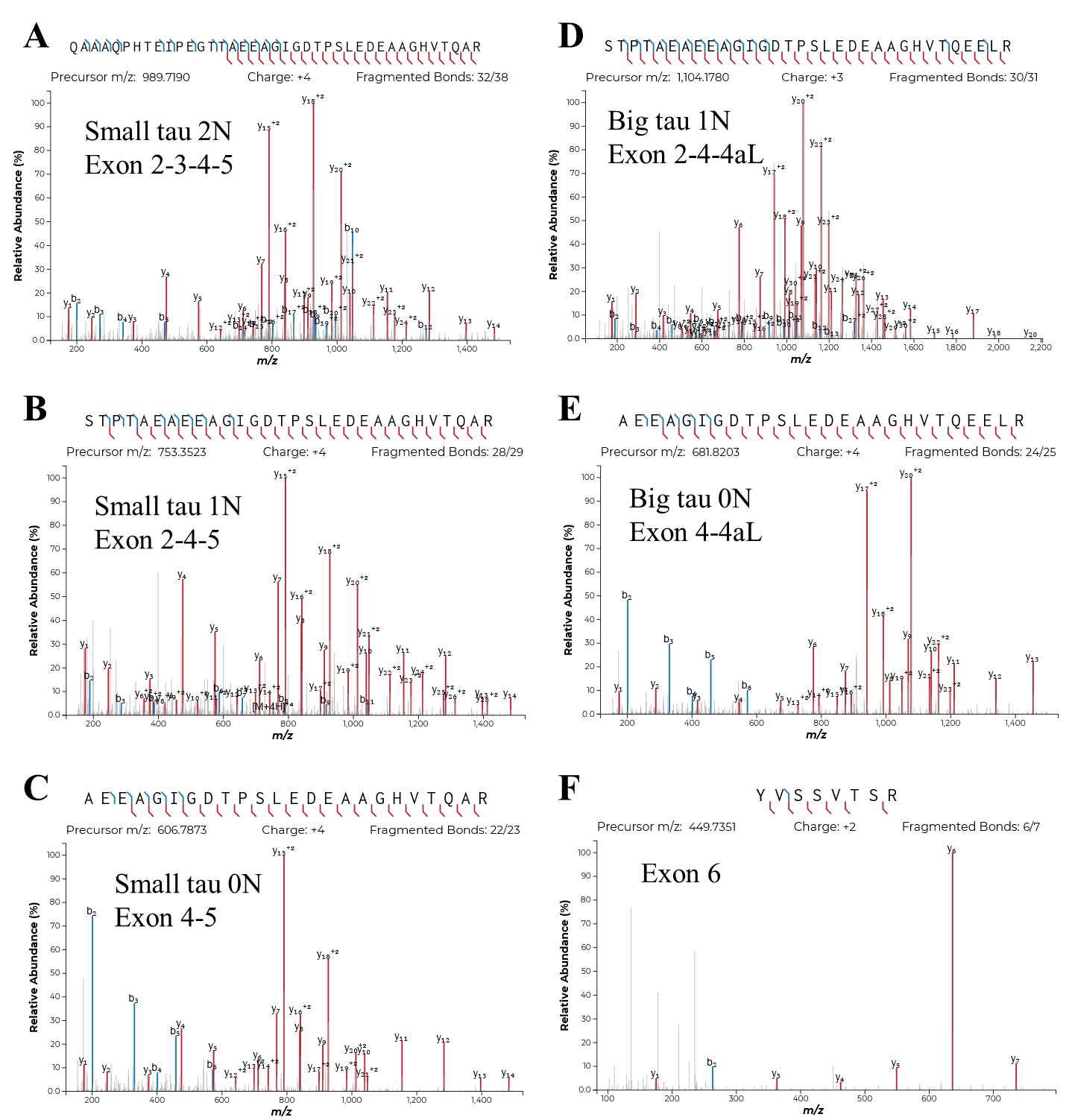
**Supplementary Figure 2**. Tandem mass spectrometry of small and big tau isoform specific peptides. Illustrative HCD-MS/MS (using NCE 30) spectra of (A) [M+4H]^4+^ *m/z* 989.7190, 2N small tau; (B) [M+4H]^4+^ *m/z* 753.3523, small tau 1N; (C) [M+4H]^4+^ *m/z* 606.7873, small tau 0N; (D) [M+3H]^3+^ *m/z* 1104.1780, big tau 1N and (E) [M+4H]^4+^ *m/z* 681.8203, big tau 0N peptides and (F) [M+2H]^2+^ *m/z* 449.7351 peptide from tau exon 6. The *y* and *b* ions are shown in red and blue, respectively. (continued in next page)

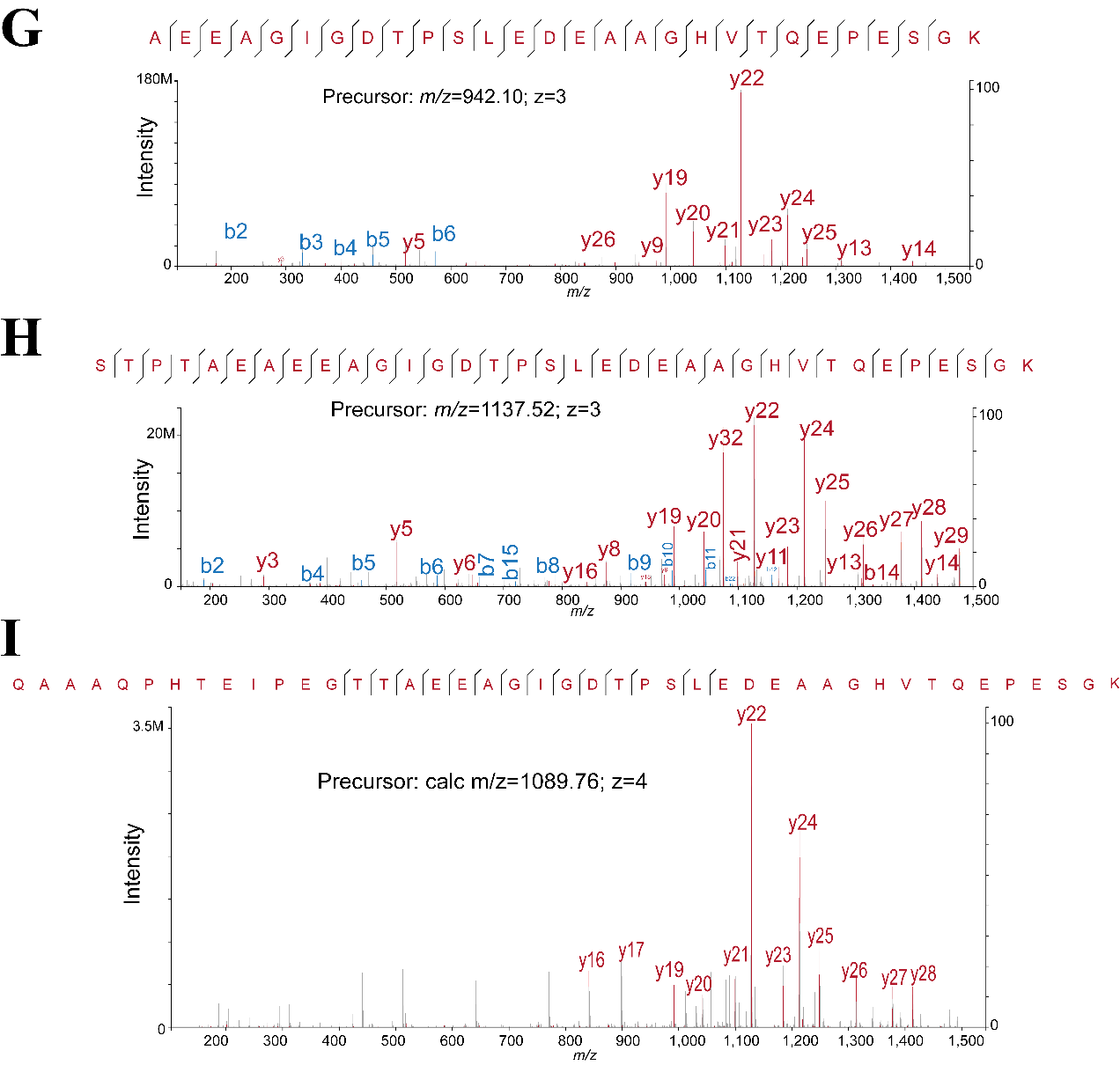
**Supplementary Figure 2 (continued)**. Tandem mass spectrometry of tau exon 4 to exon 4-S, isoform specific junction peptides (synthetic standard). Illustrative HCD-MS/MS (using NCE 30) spectra of (G) [M+3H]^3+^ *m/z* 942.0967, Big tau 0N-4a-S; (H) [M+3H]^3+^ *m/z* 1137.5166, Big tau 1N-4a-S; and (I) [M+4H]^4+^ *m/z* 1089.7560, Big tau 2N-4a-S. NCE, normalized collision energy.

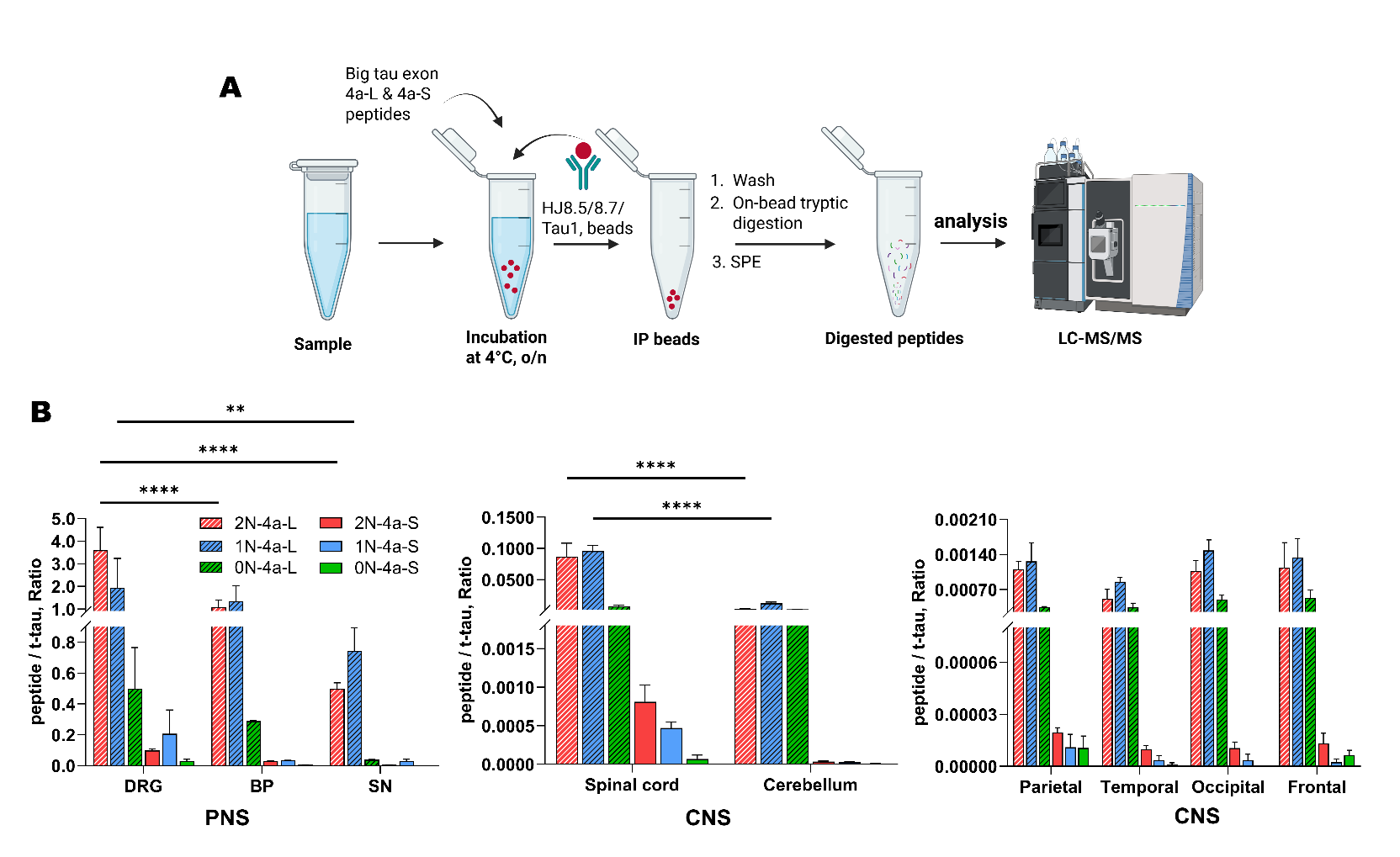

**Supplementary Figure** **3**. (A) Schematic overview of the immunoprecipitation (IP)-LC-MS/MS workflow used for the detection of exon 4a-L and exon 4a-S, big tau specific junction peptides from human nervous tissue using anti-tau antibody cocktail (HJ8.5, HJ8.7 and Tau 1) conjugated to sepharose beads and compared with the respective synthetic standard peptides. (B) Quantification of exon 4a-L and exon 4a-S, big tau peptides containing different N-terminal insert isoforms (2N, 1N, 0N) in PNS (DRG, n=2, BP, n=2, SN, n=5) and CNS (parietal, temporal, occipital, frontal, cerebellum, spinal cord, each n=6) tissue soluble lysate samples. Peptide abundances are expressed as ratios normalized to total (^212^TPSLPTPPTR^221^). Bars represent mean ± SEM. Statistical significance is indicated as *****p* < 0.0001, ***p* = 0.0082

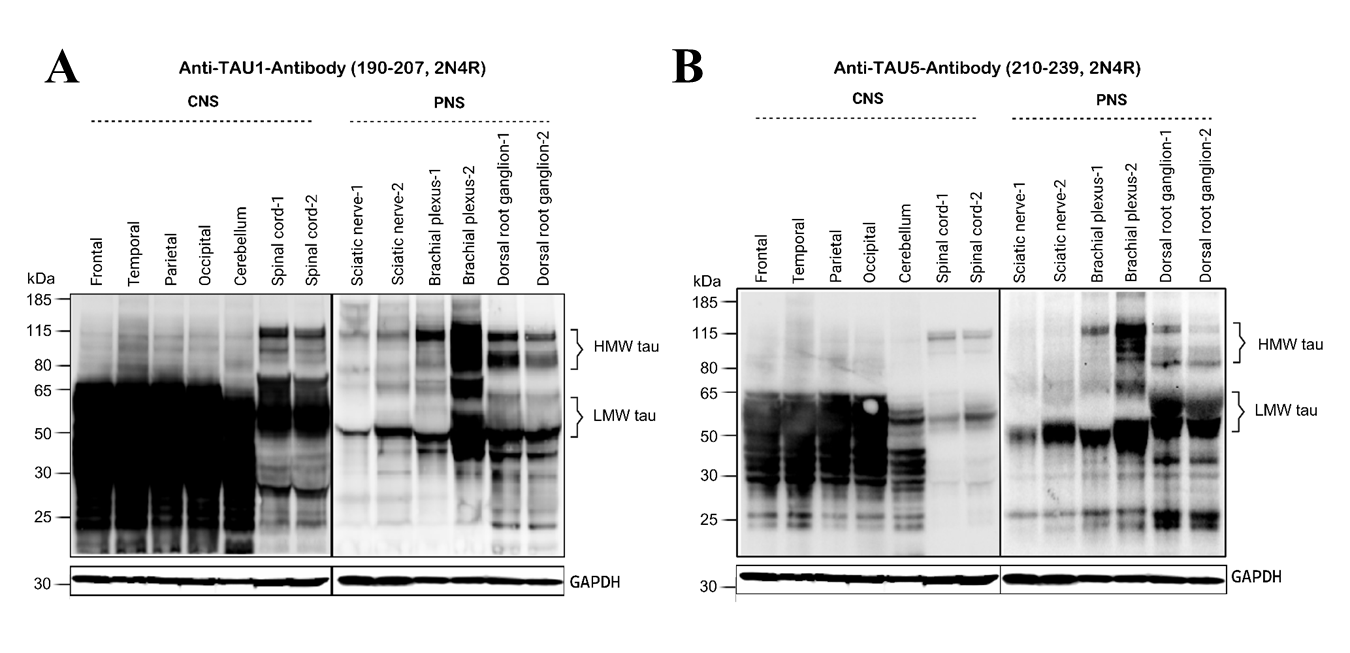
**Supplementary Figure 4.** Western blot analysis of tau protein isoforms across central and peripheral nervous tissues in humans. Representative immunoblot of (a) Tau-1 (190-207, 2N4R) and (b) Tau-5 (210-239, 2N4R) antibodies showing the expression of big tau (~115-90 kDa) and small tau isoforms (~40-65 kDa) peripheral tissues (sciatic nerve, brachial plexus, dorsal root ganglion) and spinal cord tissues. No discernable HMW bands were detected in the brain regions (temporal, parietal, occipital, frontal cortex, cerebellum) using either Tau-1 and Tau-5 antibodies. GAPDH (35 kDa) was used as loading control.

**Supplementary Figure 5.** SDS-PAGE gel stained with (A) Oriole (BioRad) fluorescent stain to visualize protein bands from brain, spinal cord, cauda equina (CE), brachial plexus (BP), and dorsal root ganglion (DRG) tissue lysates. A schematic diagram illustrates the processing for in-gel digestion and LC-MS/MS analysis of the gel bands (HMW and LMW). Gel bands corresponding to big tau (75–185 kDa) and small tau (40-70 kDa) were excised for downstream analysis. LC-MS/MS analysis was performed on peptides excised from brain (frontal), spinal cord (cervical), cauda equina (CE), brachial plexus (BP), Dorsal root ganglion (DRG) fractions. (B) Shown representative chromatogram displays extracted ion chromatograms (xic) for the major fragments of the targeted endogenous (^14^N) peptides of big tau-2N, 1N, 4R:720-738, 3R:727-738 from excised gel bands between 75-185 kDa. (C) The data represent qualitative estimations of big tau-exon 4a-L isoforms (abundance) across nervous tissue. Big tau-2N-4a-L and 1N-4a-L isoforms show the strongest to moderate levels and big tau 0N-4a-L isoform detectable only at low levels across nervous tissue.

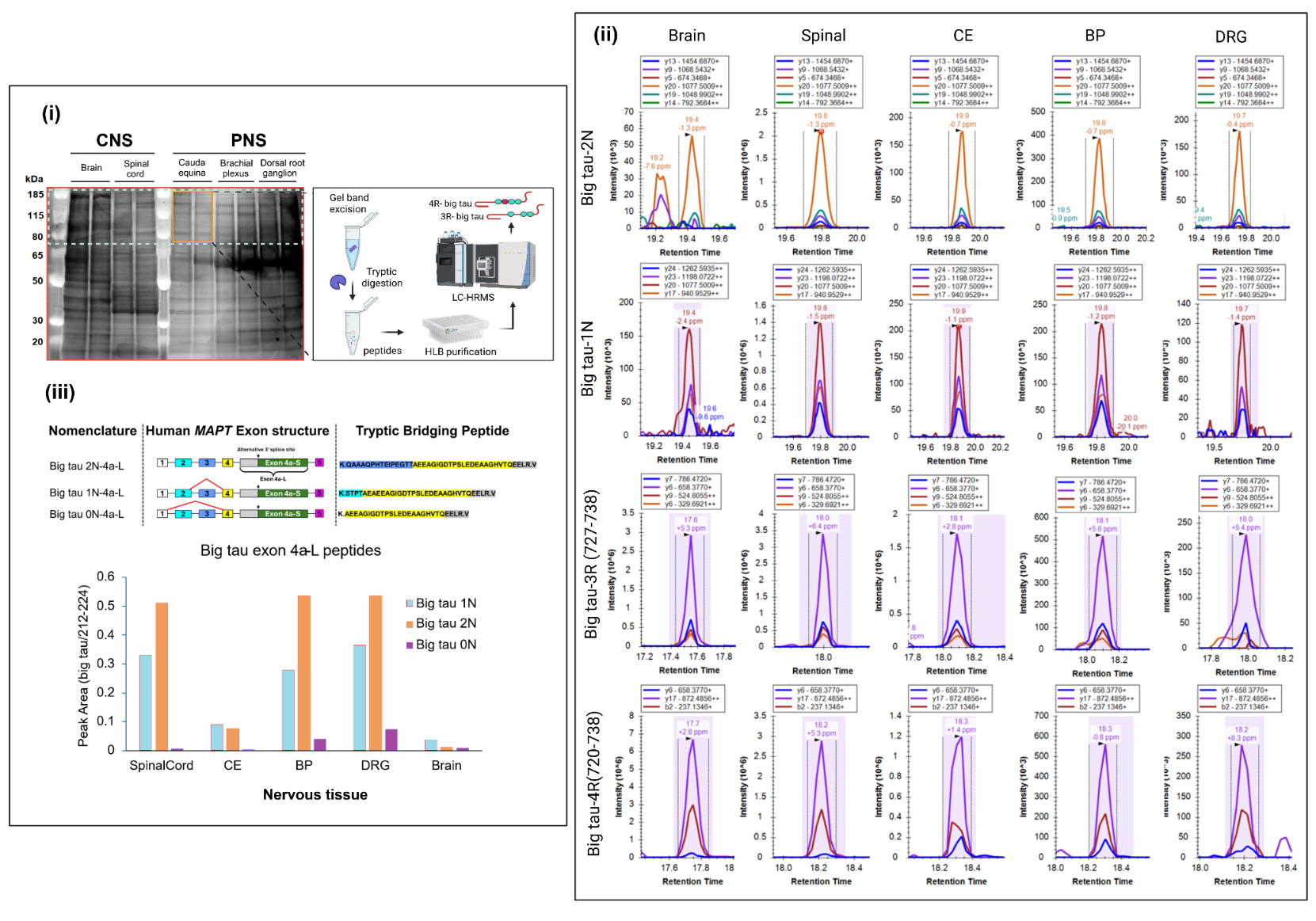

**A**

**B**

**C**

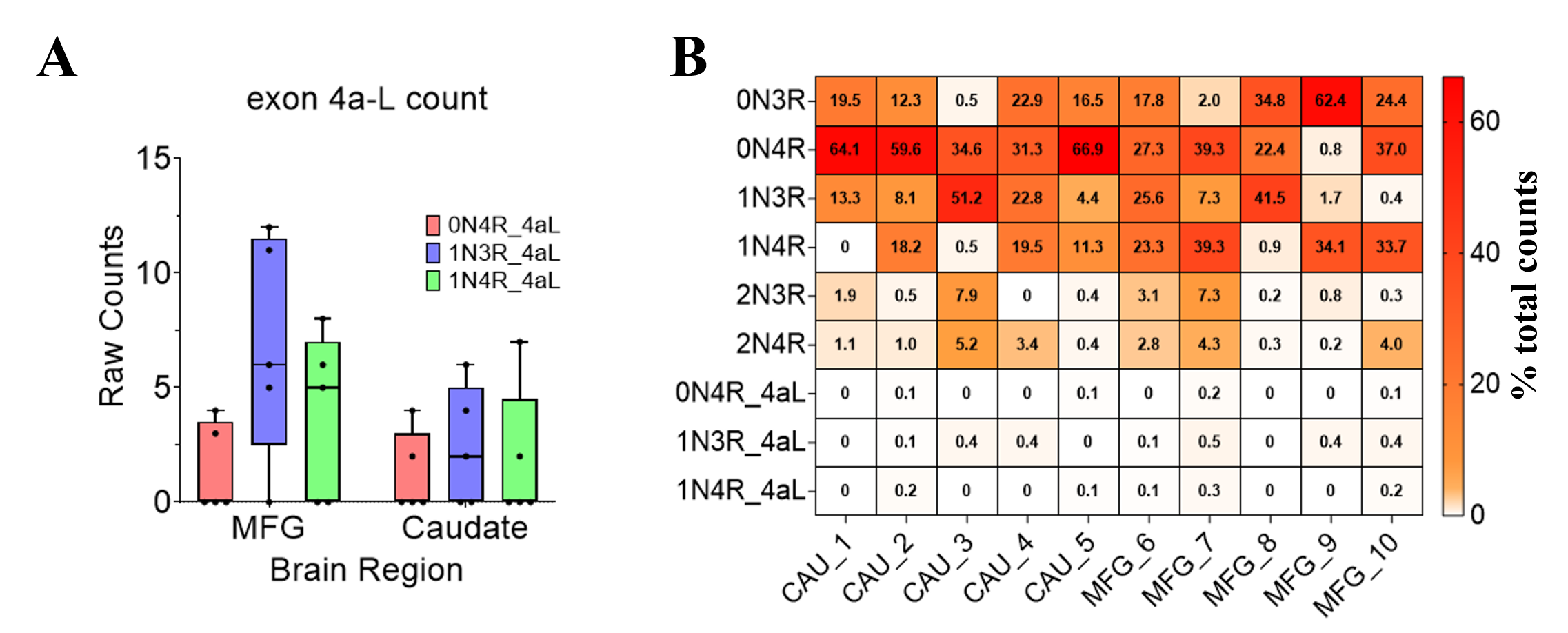

**Supplementary Figure 6:** Big tau transcripts expression in the brain. A) Bar-plot of read counts for all detectable big tau transcripts (0N3R4aL, 1N3R4aL, 1N4R4aL) in frontal cortex (MFG) and caudate (CAU), each brain is represented by a dot, only brains with at least one big tau transcript are shown. B) Heatmap of the six canonical isoforms (small tau) and the three detected big tau isoforms as percentage expression to total sample count in all the brain tissue samples investigated for transcripts.

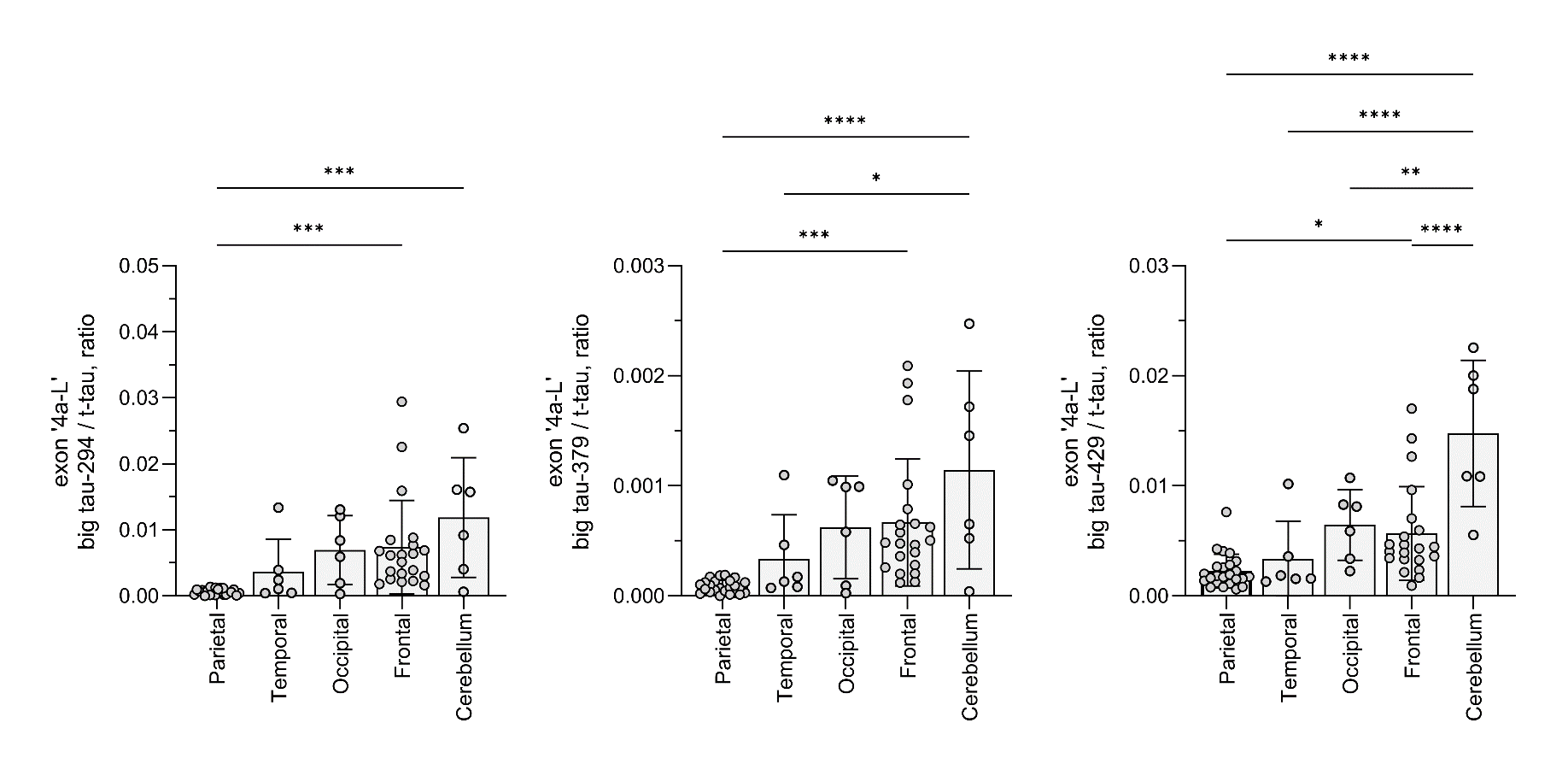
**Supplementary Figure 7.** Bar graph of relative ratio of tau exon 4a-L peptides to total tau peptide (residue 633-642) in five different regions of the adult human brain. Cerebellum (n=6) contains significantly higher compared to other cortical regions (parietal, temporal, occipital, frontal cortex) of the brain. One-Way ANOVA statistical analysis with multiple comparisons test using Dunnett statistics, ****p* < 0.0001, ***p* ≤ 0.0058, **p* ≤ 0.0432.

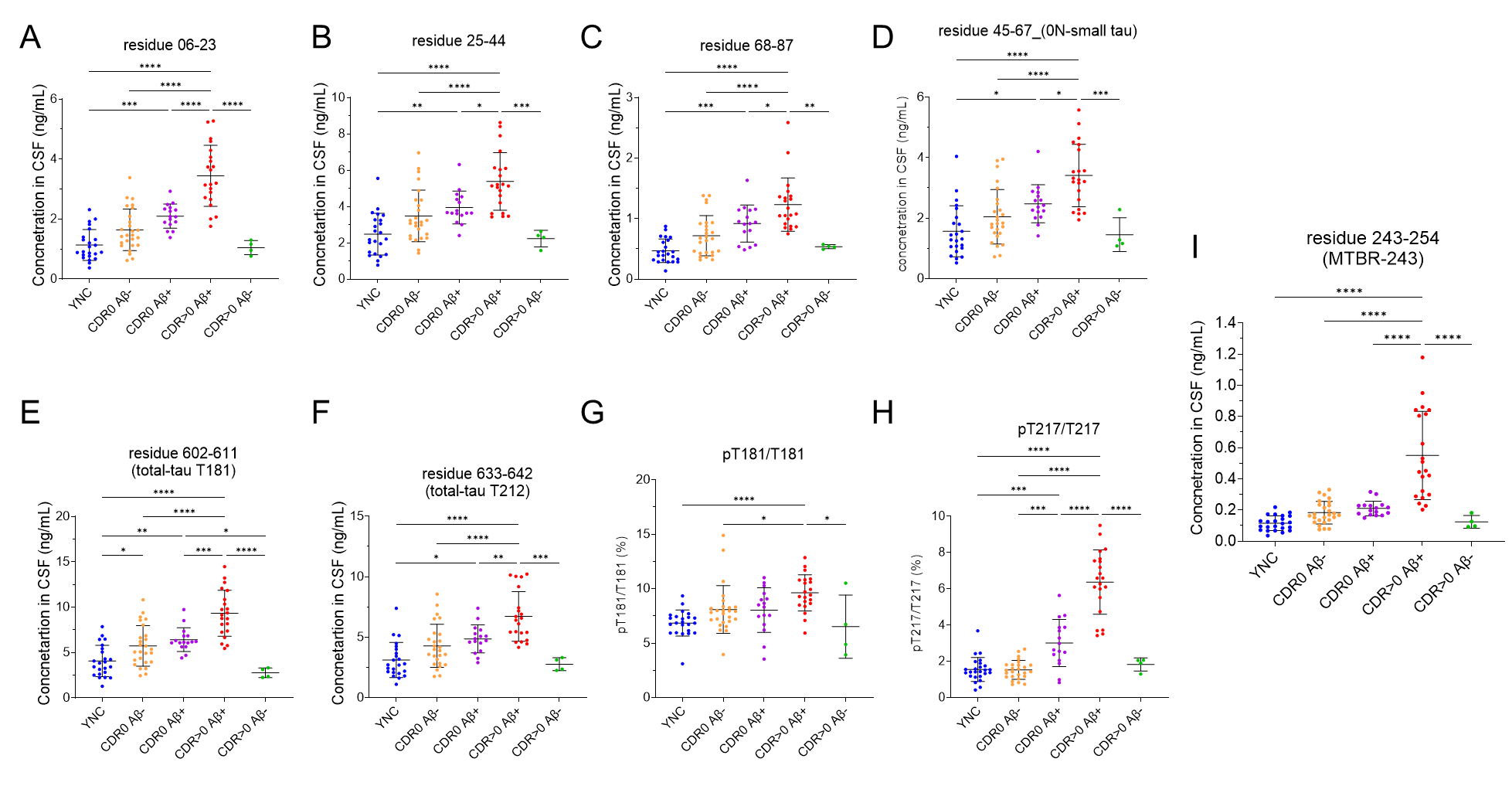

**Supplementary Figure** **8.** Scatter plots of concentration (ng/mL) in CSF of (A) N-terminal tau (residue 06-23), (B) residue 25-44, (C) residue 68-87, (D) residue 45-67 (small tau 0N), (E) residue 602-611 (total tau T181), (F) residue 633-642 (total-tau T212), (G) pT181/T181 (%), (H) pT217/T217 (%) and (I) MTBR-tau243 tau peptides from young normal controls (YNC), asymptomatic (CDR=0) Aβ-, asymptomatic (CDR=0) Aβ+, symptomatic (CDR=0) Aβ- and symptomatic (CDR=0) Aβ+ individuals. The levels of N-terminal tau-6, total tau-181, total tau-212 and MTBR-tau243 are significantly increased in the symptomatic stages of the amyloidosis (CDR=0, Aβ+) compared to asymptomatic stage (CDR>0, Aβ+) that correlates with the disease progression in AD. One-Way ANOVA with multiple comparisons test using Tukey test, ****p* < 0.0001, ***p* ≤ 0.01, **p* ≤ 0.05

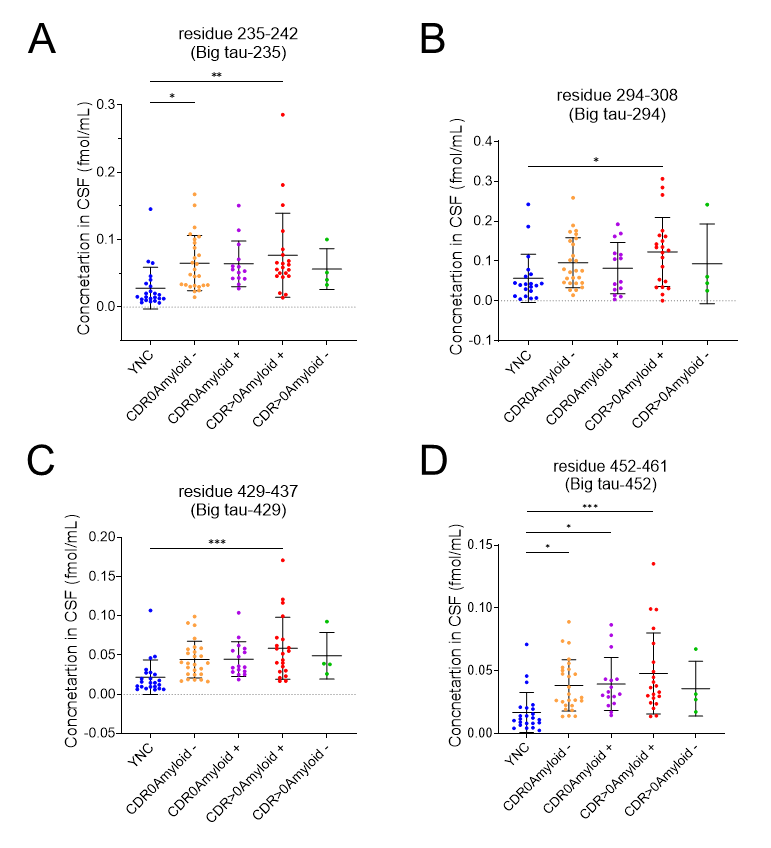
**Supplementary Figure 9.** Scatter plots for (A) big tau-235 (fmol/mL), (B) big tau-294 (fmol/mL), (C) big tau-429 (fmol/mL) and (D) big tau-452 (fmol/mL) in the CSF of young normal controls (YNC), asymptomatic (CDR=0) Aβ-, asymptomatic (CDR=0) Aβ+, symptomatic (CDR=0) Aβ- and symptomatic (CDR=0) Aβ+ individuals. One-Way ANOVA with multiple comparisons test using Tukey test, ****p* < 0.0001, ***p* ≤ 0.01, **p* ≤ 0.05

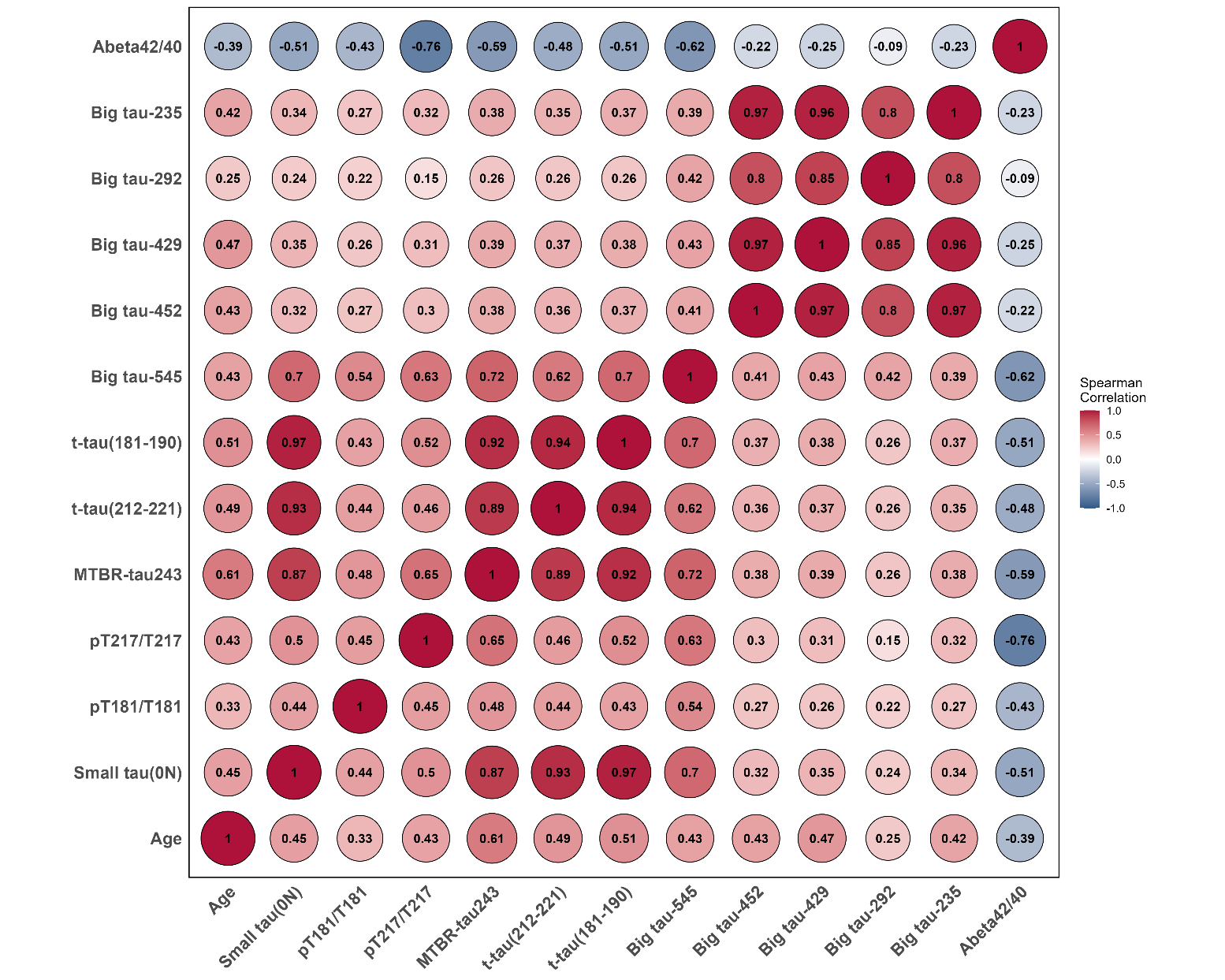
**Supplementary Figure 10:** Pair-wise Spearman’s rank correlation (ρ) of the CSF biomarkers reveals while exon 4a-L peptides correlate highly with each other, exon 6 peptide correlates with MTBR-tau243. The MTBR-tau243 correlated highly with the total tau peptides (T181 and T212) and less so with the pT217/T217 and T181/T181. Most interestingly, we observed the exon ‘4a-L’ and ‘6’ tau peptides correlate with age of the individuals, just like MTBR-tau243 and total tau in the CSF cohort that includes young normal controls, Aβ- and Aβ+ individuals.

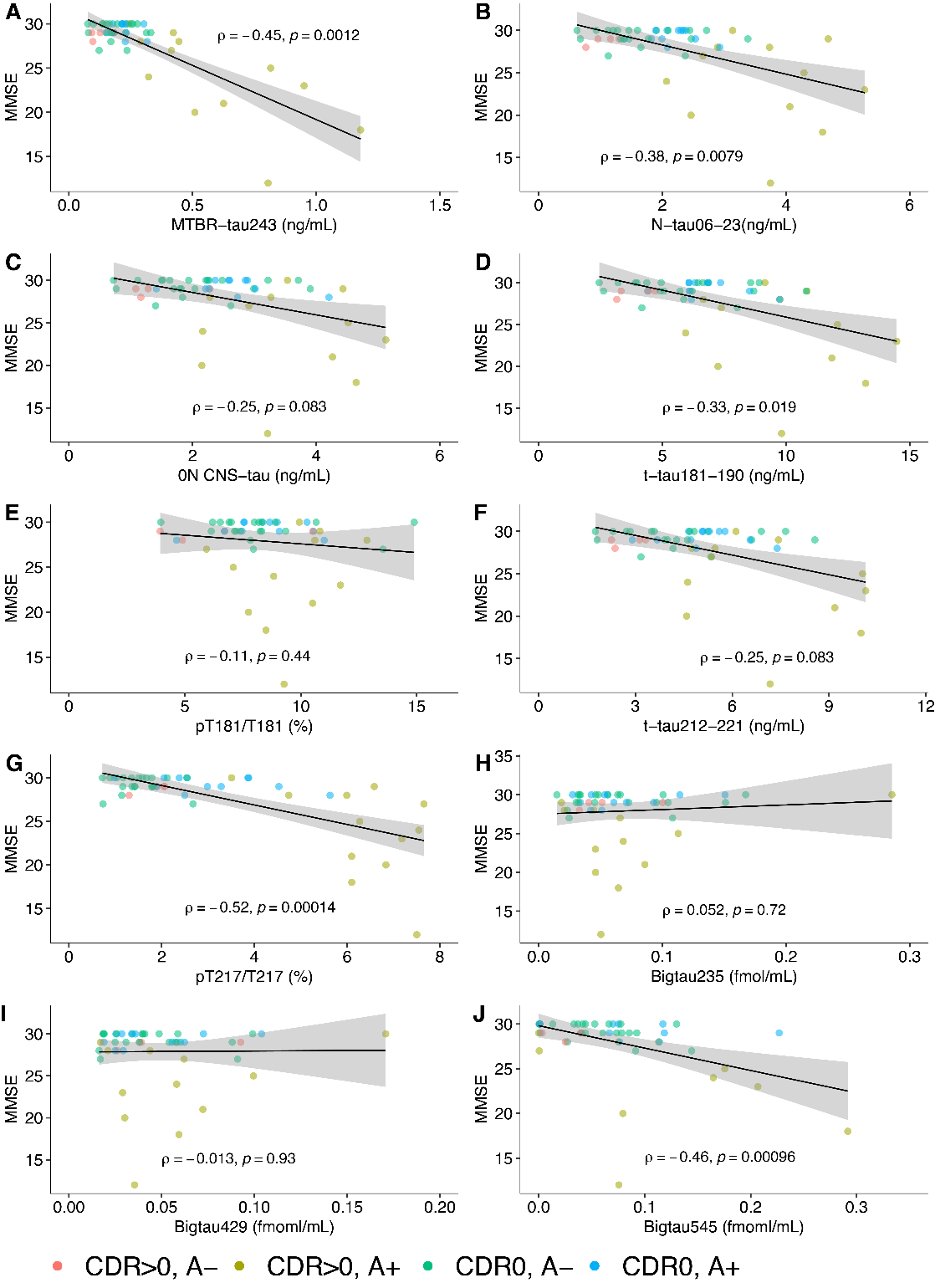

**Supplementary Figure 11**. **Associations between CSF biomakers and MMSE**. Scatter plots depicting the association between (A) MTBR-tau243 (ng/mL), (B) N-terminus tau06-23 (ng/mL), (C) CNS 0N specific tau (ng/mL), (D) pT181/T181 occupancy (%) , (E) pT217/T217 occupancy (%), (F) t-tau181-190 (ng/mL) and (G) t-tau212-221 (ng/mL), 9H) Bigtau235 (fmol/mL), (I) Bigtau429 (fmol/mL) and MMSE in the CSF cohort, color-coded by clinical diagnosis (CDR) and amyloid status (A-/A+).

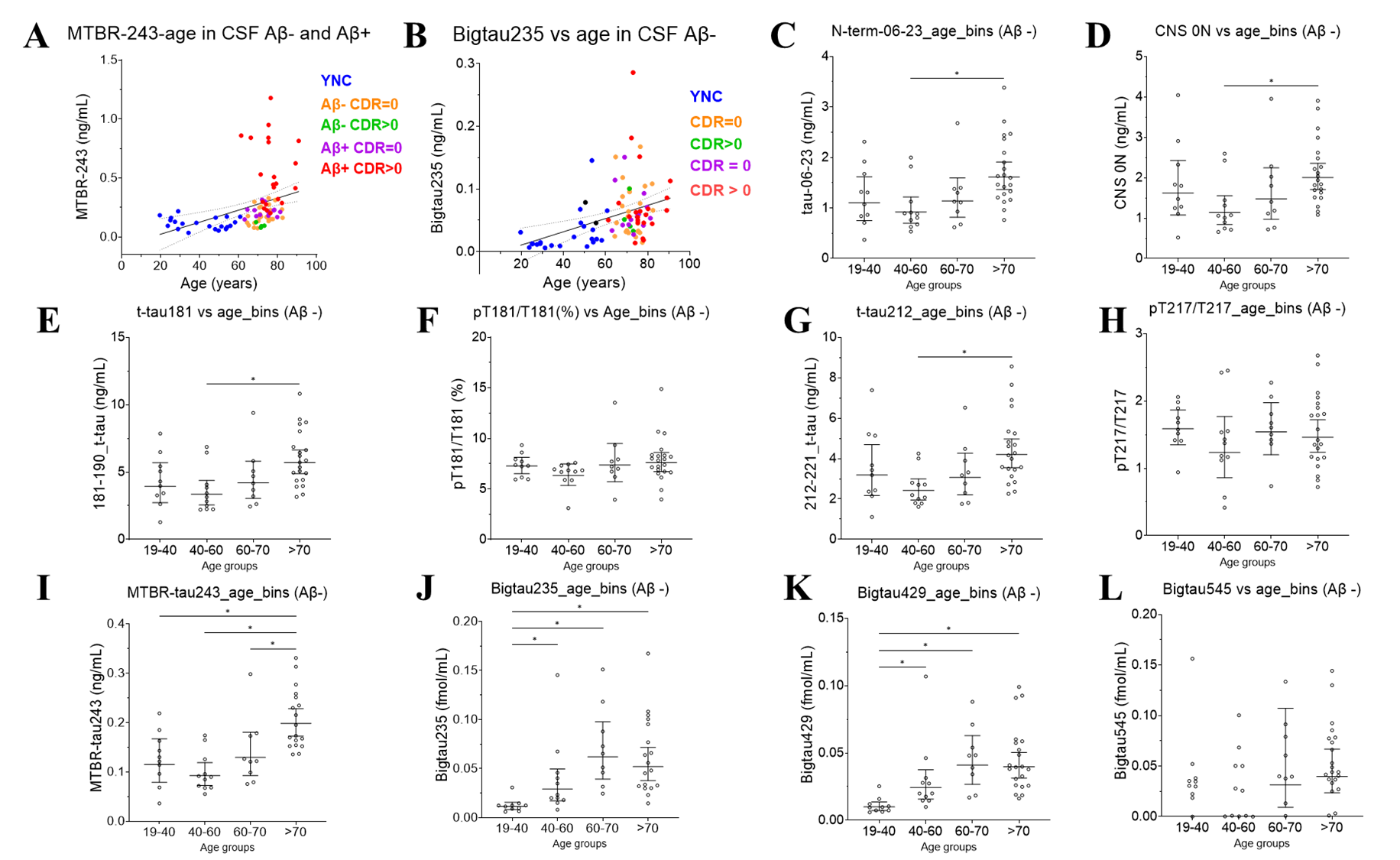
**Supplementary Figure 12.** Association of CSF biomarkers with age in Aβ+ and Aβ- individuals. (A) MTBR-tau243 (ng/mL), (B) Bigtau235 (fmol/mL) in both all Aβ+ and Aβ- individuals. The CSF biomarkers increase in Aβ- individuals with different age groups. (C) N-terminus tau06-23 (ng/mL), (D) CNS 0N specific tau (ng/mL), (E) t-tau181-190 (ng/mL), (F) pT181/T181 occupancy (%), (G) t-tau212-221 (ng/mL), (H) pT217/T217 occupancy (%), (I) MTBR-tau243 (ng/mL), (J) Bigtau235 (fmol/mL), (K) Bigtau429 (fmol/mL) and (L) Bigtau545 (fmol/mL). Statistical analysis was performed with Kruskal-Wallis test with false discovery rate correction for multiple comparison with Benjamini-Hochberg method.

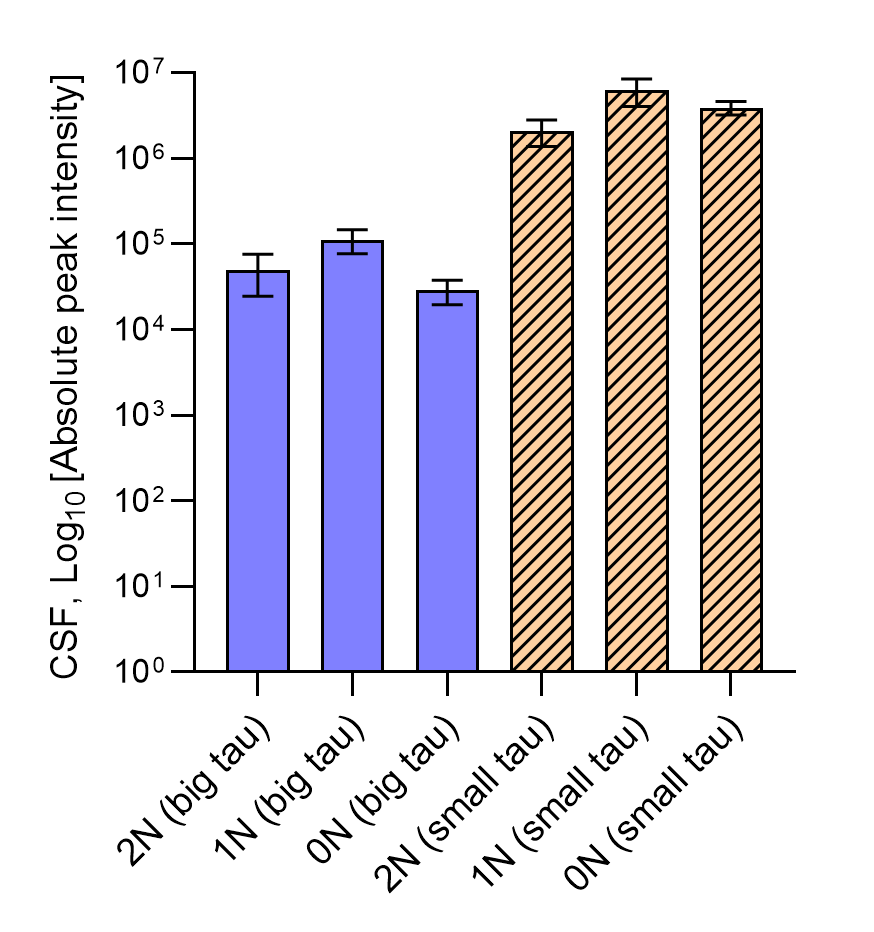

**Supplementary Figure 13.** Exon 4a-L big tau N-insert isoforms (2N, 1N, 0N) were detected and quantified in cerebrospinal fluid (CSF) alongside small tau N-insert isoforms. The relative abundance of big tau N-inserts in CSF was approximately two-orders of magnitude lower than small tau counterparts.

| **Target peptide sequence** | **Residue** | **Exon location on *MAPT*** | **Digestion enzyme(s)** | **MS Methodology** |
| --- | --- | --- | --- | --- |
| STPTAEAEEAGIGDTPSLEDEAAGHVTQAR – 1N (CNS) | 68-97 | Exon 2 and 4; exclusion of Exon 3 | Trypsin | PRM/DDA |
| STPTAEAEEAGIGDTPSLEDEAAGHVTQEELR -1N (PNS) | 68-99 | Exon 2, 4 and 4a-L; exclusion of Exon 3 | Trypsin | PRM/DDA |
| QAAAQPHTEIPEGTTAEEAGIGDTPSLEDEAAGHVTQAR – 2N (CNS) | 88-126 | Exon 3 to 4 | Trypsin | PRM/DDA |
| QAAAQPHTEIPEGTTAEEAGIGDTPSLEDEAAGHVTQEELR – 2N (PNS) | 88-128 | Exon 3, 4 and 4a-L | Trypsin | PRM/DDA |
| AEEAGIGDTPSLEDEAAGHVTQAR – 0N (CNS) | 45-68 | Exon 4 | Trypsin | PRM/DDA |
| AEEAGIGDTPSLEDEAAGHVTQEELR – 0N (PNS) | 45-70 | Exon 4 to 4a-L | Trypsin | PRM/DDA |
| DEAAGHVTQAR (CNS) | 116-126 | Exon 4 | Asp-N + Trypsin | PRM/DDA |
| DEAAGHVTQEELR (PNS) | 116-128 | Exon 4 to 4a-L | Asp-N + Trypsin | PRM/DDA |
| APERPLANEISAHVQPGPCGEASGVSGPCLGEK | 136-168 | Exon 4a-L | Asp-N + Trypsin | PRM |
| ISAHVQPGPCGE | 145-156 | Exon 4a-L | Glu-C + Trypsin | PRM |
| EPEAPVPLTASLPQHRPVCPAPPPTGGPQEPSLEWGQK | 169-206 | Exon 4a-L | Asp-N + Trypsin | PRM |
| GPAFPKPATTAYLHTEPESGK | 214-234 | Exon 4a-L | Asp-N + Trypsin | PRM/DDA |
| VVQEGFLR | 235-242 | Exon 4a-L | Trypsin | PRM/DDA |
| EPGPPGLSHQLMSGMPGAPLLPEGPR | 243-268 | Exon 4a-L | Asp-N + Trypsin | PRM |
| HQLLGDLHQEGPPLK | 294-308 | Exon 4a-L | Trypsin | PRM/DDA |
| VSTEIPASEPDGPSVGR | 379-395 | Exon 4a-L | Trypsin | PRM/DDA |
| AAFPGAPGEGPEAR | 429-442 | Exon 4a-L | Trypsin | PRM/DDA |
| EADLPEPSEK | 452-461 | Exon 4a-L | Trypsin | PRM/DDA |
| HPTPGSSDPLIQPSSPAVCPEPPSSPK | 518-544 | Exon 6 | Trypsin | PRM |
| YVSSVTSR | 545-552 | Exon 6 | Trypsin | PRM/DDA |

**Supplementary Table 1** The list of peptides identified using multi-enzyme digest. C, carbamidomethyl.

**Supplementary Table 2**. The list of synthetic peptides from exon 4-exon 4a-L and exon 4-exon 4a-S bridging regions and their corresponding tryptic peptide used for validation in this study

| Nomenclature | Synthetic Peptide Sequence | Residue | Tryptic Bridging Peptide |
| --- | --- | --- | --- |
| 1N Big Tau E4a-L | STPTAEAEEAGIGDTPSLEDEAAGHVTQEELRVPGRQR | 68-105 | STPTAEAEEAGIGDTPSLEDEAAGHVTQEELR |
| 2N Big Tau E4a-L | QAAAQPHTEIPEGTTAEEAGIGDTPSLEDEAAGHVTQEELRVPGRQR | 88-134 | QAAAQPHTEIPEGTTAEEAGIGDTPSLEDEAAGHVTQEELR |
| 0N Big Tau E4a-L | AEEAGIGDTPSLEDEAAGHVTQEELRVPGRQR | 45-76 | AEEAGIGDTPSLEDEAAGHVTQEELR |
| 1N Big Tau E4a-S | STPTAEAEEAGIGDTPSLEDEAAGHVTQEPESGKVVQEGFLR | 68-109 | STPTAEAEEAGIGDTPSLEDEAAGHVTQEPESGK |
| 2N Big Tau E4a-S | QAAAQPHTEIPEGTTAEEAGIGDTPSLEDEAAGHVTQEPESGKVVQEGFLR | 88-138 | QAAAQPHTEIPEGTTAEEAGIGDTPSLEDEAAGHVTQEPESGK |
| 0N Big Tau E4a-S | AEEAGIGDTPSLEDEAAGHVTQEPESGKVVQEGFLR | 45-80 | AEEAGIGDTPSLEDEAAGHVTQEPESGK |

**Supplementary Table 3.** Clinical demographics of brain tissues used for MAPT Targeted isoseq. Abbreviations: CAU, caudate; MFG, middle frontal gyrus; RIN, RNA Integrity Number; F, female; M, male; PMI, postmortem interval.

| **Brain_ID** | **Brain region** | **Sample_ID** | **Brain_Bank** | **sex** | **RIN** | **PMI (hours)** | **Age at death (years)** |
| --- | --- | --- | --- | --- | --- | --- | --- |
| CAU_1 | CAU | 6904-cau | UWA | F | 6.1 | 5.9 | 83 |
| CAU_2 | CAU | 2020-054-2-cau-put | NBB | F | 5 | 0 | 81 |
| CAU_3 | CAU | HBCC-1987-cau | HBCC | M | 8.8 | 27 | 54 |
| CAU_4 | CAU | HBCC-2871-cau | HBCC | M | 8.9 | 20 | 61 |
| CAU_5 | CAU | NBB-2019-116-cau | NBB | M | 5 | 0 | 79 |
| MFG_6 | MFG | BSHRI-2015-1433-MFG | BSHRI | M | 4.3 | 2 | 63 |
| MFG_7 | MFG | BSHRI-2015-280-MFG | BSHRI | F | 7.2 | 4.2 | 63 |
| MFG_8 | MFG | BSHRI-2015-735-MFG | BSHRI | M | 5.1 | 4.2 | 63 |
| MFG_9 | MFG | BSHRI-2015-1431-MFG | BSHRI | F | 7.2 | 2.5 | 77 |
| MFG_10 | MFG | BSHRI-2015-1401-MFG | BSHRI | F | 6.2 | 1.5 | 82 |
